# BiOS: An Open-Source Framework for the Integration of Heterogeneous Biodiversity Data

**DOI:** 10.64898/2026.02.09.704853

**Authors:** Alejandro Roldán, Tomás Golomb Durán, Antoni Josep Far, Maria Capa, Enrique Arboleda, Tommaso Cancellario

## Abstract

The era of Big Data has reshaped biodiversity research, yet the potential of this information is frequently constrained by data heterogeneity, incompatible schemas, and the fragmentation of resources. Whilst standards such as Darwin Core have improved interoperability, significant barriers persist in harmonising multi-typology datasets ranging from taxonomy and genetics to species distribution. Here, we present the Biodiversity Observatory System (BiOS), a comprehensive, open-source software stack designed to address these impediments through a modular, community-driven architecture.

BiOS departs from monolithic database designs by decoupling the back-end data management from the front-end presentation layer. This architectural separation supports a dual-access model tailored to diverse stakeholder needs. For researchers and developers, the system offers a comprehensive Application Programming Interface (API) that exposes all back-end functionalities, enabling seamless programmatic access, automated data retrieval, and integration with external analytical workflows. Simultaneously, the platform features a user web interface designed to lower the technical barrier to entry. This interface facilitates intuitive data exploration through agile taxonomic navigation, advanced geospatial map viewers for species occurrence filtering, and dedicated dashboards for visualising genetic markers and legislative status.

Strictly adhering to the FAIR principles (Findable, Accessible, Interoperable, Reusable), BiOS acts as a relational engine capable of integrating heterogeneous data streams. By providing a flexible, interoperable core that supports the “seven shortfalls” framework of biodiversity knowledge, BiOS offers a turnkey solution to overcome data fragmentation and enhance collaborative conservation efforts.

## 1. Introduction

Biodiversity research has fully entered the era of Big Data, providing access to vast amounts of information on an unprecedented temporal and spatial scales. This innovation has facilitated data exchange and improved collaboration among researchers and institutions worldwide. Nowadays, a wide range of biodiversity records is available through public databases (Hobern et al., 2019), supporting numerous research initiatives and enhancing nature conservation efforts. These virtual resources include long-term monitoring programmes as the International Long Term Ecological Research (ILTER; Vanderbilt & Gaiser, 2017), genetic data collections (e.g., the Barcode of Life Data Systems, BOLD; Ratnasingham & Hebert, 2007), data aggregators (e.g., the Global Biodiversity Information Facility, GBIF; GBIF.org, 2025), taxonomic checklists (e.g., the Catalogue of Life, COL; Bánki et al., 2025), and trait databases (e.g., the TRY Plant Trait Database, TRY; Kattge et al., 2020). All together, these platforms host billions of records (e.g., 2.2 million accepted species in COL; 15 million plant trait records in the TRY; over 3.5 billion occurrence records in GBIF; more than 1 billion BOLD; all accessed 22 December 2025), opening the door to the investigation of ecological questions that were previously intractable due to limitations in data quality and availability. Moreover, this data availability may facilitate addressing the “seven shortfalls” described by Hortal et al. (2015), which highlight gaps in taxonomic, geographic, and ecological knowledge.

Despite the exponential growth of biodiversity information, significant barriers still hinder the effective integration and full exploitation of these data. Among these impediments are the limited awareness of existing databases and their contents, and the heterogeneity of data and database schemas (see Feng et al., 2025 for a detailed review). For example, incompatibilities among backbone taxonomic systems represent a long-recognised challenge that substantially hinders interoperability between biodiversity databases (Grenié et al., 2023; Sandall et al., 2023; Feng et al., 2022). Indeed, biological databases may rely on different taxonomic concepts (e.g., morphological *vs* molecular identification) or differ in taxonomic breadth (e.g., taxon-specific *vs* multi-taxon coverage), leading to the same biological entities being misassigned to different taxa across data sources. Moreover, the lack of regular taxonomic updates further contributes to inconsistencies, as outdated nomenclature may persist, hindering large-scale data synthesis (Grenié et al., 2023). Additional challenges arise from the heterogeneous schemas used to structure these databases (e.g., Atlas of Living Australia *vs* Global Biodiversity Information Facility; Mesibov, 2018) which often reflect traditional methodologies, discipline-specific terminology, or the conventions of individual researchers and institutions. The challenge is compounded by differences in the underlying informatics systems, such as back-end structures and data management frameworks, which are often incompatible across platforms. Together, these impediments make it difficult to create harmonised and comprehensive datasets, leading researchers to spend considerable time and effort searching, organising, reconciling taxonomies, and normalising data, rather than focusing on answering scientific questions.

To address some of the issues described, standards and protocols such as Darwin Core (Wieczorek et al., 2012) and the Veg-X protocol (Wiser et al., 2011) have been developed and have demonstrated that their widespread adoption and structured schemas can significantly enhance the usability and interoperability of biodiversity data. These frameworks provide essential guidelines and standardised vocabularies for recording taxonomic, spatial, temporal, and ecological information, enabling more effective data sharing and reuse across platforms and research communities. In practice, their implementation has facilitated the aggregation of datasets from multiple sources, supported large-scale biodiversity assessments, and improved reproducibility in ecological research. Despite these advances, however, only a few initiatives have achieved high levels of standardisation, and substantial heterogeneity persists across databases and resources. Differences in data formats, metadata completeness, and terminologies continue to limit the seamless integration of biodiversity datasets, hindering large-scale analyses and cross-institutional collaborations. Moreover, many existing tools focus on specific data types or taxonomic groups, making it challenging to adopt a unified approach for multi-typology biodiversity data. This ongoing fragmentation underscores the need to develop more flexible, interoperable, and community-driven platforms capable of harmonising heterogeneous data whilst remaining extensible and broadly accessible.

Here, we present the foundations of a new bioinformatics tool designed to integrate and harmonise multiple types of biodiversity data, facilitating their sharing and offering potential benefits to both the scientific community and stakeholders engaged in environmental conservation. Biodiversity Observatory System (BiOS) is a relational database specifically designed to store, manage, and distribute biological and ecological data in accordance with the FAIR principles (Findable, Accessible, Interoperable, Reusable; Wilkinson et al., 2016). We aimed to develop an easily deployable and modular tool built upon a standardised core structure that allows individual database instances to be seamlessly linked. By establishing a solid structural backbone for both the back-end and the front-end architectures, this system facilitates the integration of biodiversity data from multiple sources and typologies (e.g., taxonomy, genetics, and distribution), thereby enabling more comprehensive data sharing and expanding analysis possibilities. We developed BiOS following an open-source philosophy to support a collaborative, community-driven, and modular system applicable across diverse institutional stakeholders, spanning academia, government, the non-profit sector, and industry. Rather than creating a single monolithic database, we designed BiOS to be freely deployable by users, embracing a flexible and extensible architecture. To ensure modularity and ease of maintenance, we implemented a simplified design that separates back-end components (database and API) from front-end components (web user interface). This separation allows the system to be dynamic whilst keeping all functions fully accessible through the API, enabling seamless integration with external tools and workflows. We designed the user interface to provide intuitive access for users, supporting efficient data exploration and management. By combining modular design, standardisation, and interoperability, BiOS effectively addresses current challenges in biodiversity data integration and management, facilitating large-scale analyses and promoting collaborative, community-driven research.

### 1.1 An overview of contemporary biodiversity databases

To contextualise the need for a new integrative and modular biodiversity infrastructure, we conducted a systematic review of the major public biodiversity databases, assessing both data availability and data accessibility across core biological data dimensions (Tab. 1). Our review focused on whether the system is open-source, exposes a public API, and on eight biological and ecological categories: taxonomy, georeferenced distribution, genetics, biological traits, conservation status, legislative tagging, system, and habitat. We employed a hierarchical approach to perform this review. First, we assessed whether each platform provided API documentation. If it was available and supported any of the target data categories, these categories were classified as “Yes,” indicating that the data were structured and directly accessible. In cases where no API documentation was found, we queried the platform’s website directly. For many widely studied species, data coverage in such databases is often comprehensive which helps reduce the likelihood of false negatives. Therefore, to standardise our assessment and minimise bias due to taxonomic relationships or preferences driven by the expertise of the authors, we used the species list proposed by Caldwell et al. (2024). The list comprises 27 taxa, representing the following classes: Mammalia, Reptilia, and Aves (Kingdom: Animalia) and Pinopsida, Magnoliopsida, and Polypodiopsida (Kingdom: Plantae). During the website-visiting, we determined whether the web page retrieved information via a third-party service (e.g., Wikipedia). When third-party sources were used, only English-language content was considered. If such external requests were detected, data availability was categorised as either “Yes” (i.e. data were provided in a structured format) or “Latent” (i.e. data were unstructured free-text), depending on the nature of the integration. Conversely, if no relevant data could be located through these steps, the platform was recorded as “No”. Finally, to minimise potential omissions in any category, the results were reviewed and validated by biodiversity experts.

**Table 1.** Comparison of data availability and infrastructure across major biodiversity databases. The table evaluates the presence of specific biological/ecological data categories and technical features. Data accessibility is classified into three states: Yes (green; data is structured and directly accessible); No (light blue; data is absent); Lat. for Latent (light green; data is present but unstructured, such as within free-text fields, and requires extraction). All platforms accessed 13 February 2026. Atlas of the Living Australia (ALA); BoldSystems (BOLD); Catalogue Of Life (COL); Global Biodiversity Information Facility (GBIF); INaturalist (iNat); International Union for Conservation of Nature (IUCN); National Center for Biotechnology Information (NCBI); Ocean Biodiversity Information System (OBIS); Plants Of the Word Online (POWO); World Register of Marine Species (WoRMS).

|  |  | Database |  |  |  |  |  |  |  |  |  |  |
| --- | --- | --- | --- | --- | --- | --- | --- | --- | --- | --- | --- | --- |
| Data category |  | ALA | BOLD | COL | GBIF | iNat. | IUCN | NCBI | OBIS | POWO | WoRMS | BiOS |
| Taxonomy |  | Yes | Yes | Yes | Yes | Yes | Yes | Yes | Yes | Yes | Yes | Yes |
| Georeferenced distribution |  | Yes | Yes | Lat. | Yes | Yes | Lat. | Lat. | Yes | Lat. | Lat. | Yes |
| Genetics |  | No | Yes | No | Yes | No | No | Yes | Yes | No | No | Yes |
| Biological traits<br>(e.g. body length) |  | Yes | Yes | No | Lat. | Yes | Yes | Lat. | No | Lat. | Yes | No |
| Tags | Legislative<br>(e.g., CITES) | Yes | No | No | Lat. | Yes | Yes | No | No | No | Yes | Yes |
|  | Conservation status | Yes | No | No | Yes | Yes | Yes | No | Yes | Lat. | Yes | Yes |
|  | System<br>(e.g., marine) | Lat. | Yes | Yes | Yes | No | Yes | Lat. | No | No | Yes | Yes |
|  | Habitat | Lat. | No | No | Yes | No | Yes | Lat. | No | Lat. | Yes | Yes |
| API |  | Yes | Yes | Yes | Yes | Yes | Yes | Yes | Yes | Yes | Yes | Yes |
| Open source<br>(infrastructure) |  | Yes | Yes | Yes | Yes | Yes | No | No | Yes | Yes | No | Yes |

\* Caldwell et al. (2024) species list: *Canis familiaris*; *Canis lupus*; *Ursus arctos*; *Ovis aries*; *Picea abies*; *Panthera tigris*; *Oryctolagus cuniculus*; *Caretta caretta*; *Felis catus*; *Loxodonta africana*; *Panthera pardus*; *Panthera leo*; *Chelonia mydas*; *Vulpes vulpes*; *Sus scrofa*; *Calluna vulgaris*; *Rangifer tarandus*; *Capra hircus*; *Dermochelys coriacea*; *Gallus gallus*; *Lutra lutra*; *Eretmochelys imbricata*; *Molothrus ater*; *Trichechus manatus*; *Pteridium aquilinum*; *Aythya ferina*; *Pelecanus occidentalis*.

Some considerations on the comparative assessment should be acknowledged. The evaluation was conducted using widely studied organisms to minimise taxonomic bias (Caldwell et al. 2024); however, platform functionalities evolve continuously, and the analysis reflects the state of each system at the time of access, and could eventually change in time. Furthermore, the comparison focused on the presence and structural accessibility of data categories rather than on the completeness or depth of information contained within each platform. Future work could expand the assessment to additional taxa, functional use cases, and interoperability benchmarks.

## 2. Software Tool Description

We developed BiOS using an architecture composed of three interconnected layers: i) the database and ii) the RESTful Application Programming Interface (RESTful API; thereafter API), which together compose the back-end, and iii) the web interface, which constitutes the front-end (Fig.1). We designed the architecture paying particular attention to both scalability and interoperability. These two properties are critical requirements for building a complex database designed to store large and heterogeneous biodiversity records, as they are essential for facilitating data integration from multiple sources and information exchange, which in turn enables collaboration among scientists. For a quick installation guide read *Supplementary Materials S1 & S5*.

**Figure 1.**
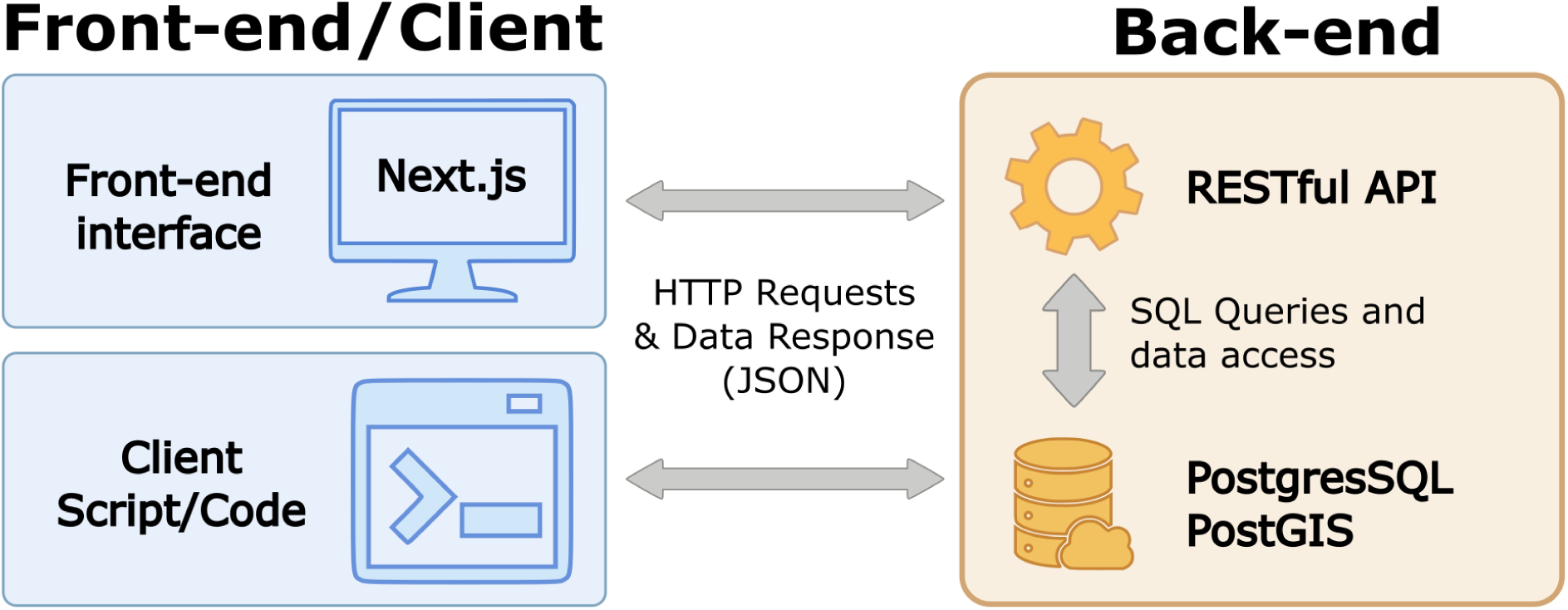
Schematic BiOS software architecture. The orange box includes the two main components of the back-end: the PostgreSQL-PostGIS database and the RESTful API service. The blue boxes represent the front-end, and the client query system. The vertical bidirectional arrow represents the interaction between the database and the API service (in the orange box), whereas the horizontal bidirectional arrows represent the HTTP request and response between the back-end and the front-end/client systems.

The back-end relies on a PostgreSQL database system (The PostgreSQL Global Development Group, 2025), and database interactions were implemented using *psycopg* (psycopg2-binary, 2022), a dedicated PostgreSQL adapter for Python (Van Rossum and Drake, 2009).

Together with PostgreSQL, we used the PostGIS extension (PostGIS Developers, 2023), which provides support for Geographic Information System (GIS) data and functions. To facilitate the database development and ensure consistent relations between tables, we implemented the entire service using Django (Django Software Foundation, 2023), a high-level Python web framework. Django is a state-of-the-art framework for developing database applications, and it offers several advantages, including a robust Object–Relational Mapping system, built-in security features, an automatically generated administrative interface for immediate data management and a clear modular structure that accelerates development workflow whilst maintaining code consistency (see Section 2.1). In addition, it offers a built-in administration platform to manage the database with a user system.

To query and interact with the database, we created a RESTful API system built using the Django REST Framework extension (Django REST Framework, 2024). The API system serves a dual purpose. First, it acts as an interface between the back-end and its users (handling both web and local client requests), allowing the web interface to present information in a user-friendly format. Second, it is essential for filtering and executing operations based on the specific parameters included by the user in their server request.

To facilitate the configuration of the database and API system, we included it in a Docker container (Docker Inc., 2025). This approach creates an isolated system for the development and production environments, ensuring efficient management of dependencies (i.e., PostGIS and Django) and increasing the reproducibility of the setup (see Section 2.2).

Finally, we developed the web interface, using Next.js 15 (Vercel Inc., 2025), due to its flexibility as this allowed us to develop a solution that was better suited to our needs (see Section 2.3).

### 2.1 Database

We organised the information contained in the database into three modules: i) thematic modules, ii) geography, and iii) versioning. This modular structure facilitates the management of diverse data types (e.g., from spatial to genetic data) whilst preserving their traceability and efficiency. It is important to note that the schema presented here provides a simplified representation of the tables and their interactions and is intended to illustrate the core concepts underlying the database design. A comprehensive and detailed description of the framework’s internal architecture, data models, and relationships (e.g., Foreign Keys, Many-to-Many fields) is reported in *Supplementary Materials S2*.

#### Thematic modules

We define a “thematic module” (structurally as a Django App), a self-contained unit responsible for a specific category of biodiversity data. Each application has a group of primary database tables designed to store homogeneous attributes (e.g., taxonomy or genetics), supported by linked auxiliary tables that provide granular detail. This design ensures a separation of concerns, maintaining a clean and organised data schema. This modular organisation facilitates data management, offering several advantages. For example, when a record within a thematic module is added or modified, the change is automatically propagated throughout the database, avoiding the need to manually review other linked modules. Furthermore, it simplifies data maintenance and scalability, allowing new datasets or functional components to be integrated without disrupting the existing architecture (for further implementation details see *Supplementary Material* S2.1).

The database is composed of six thematic modules: Taxonomy, Occurrences, Genetics, Tags, Geography, and Versioning (Fig. 2).

**Figure 2.**
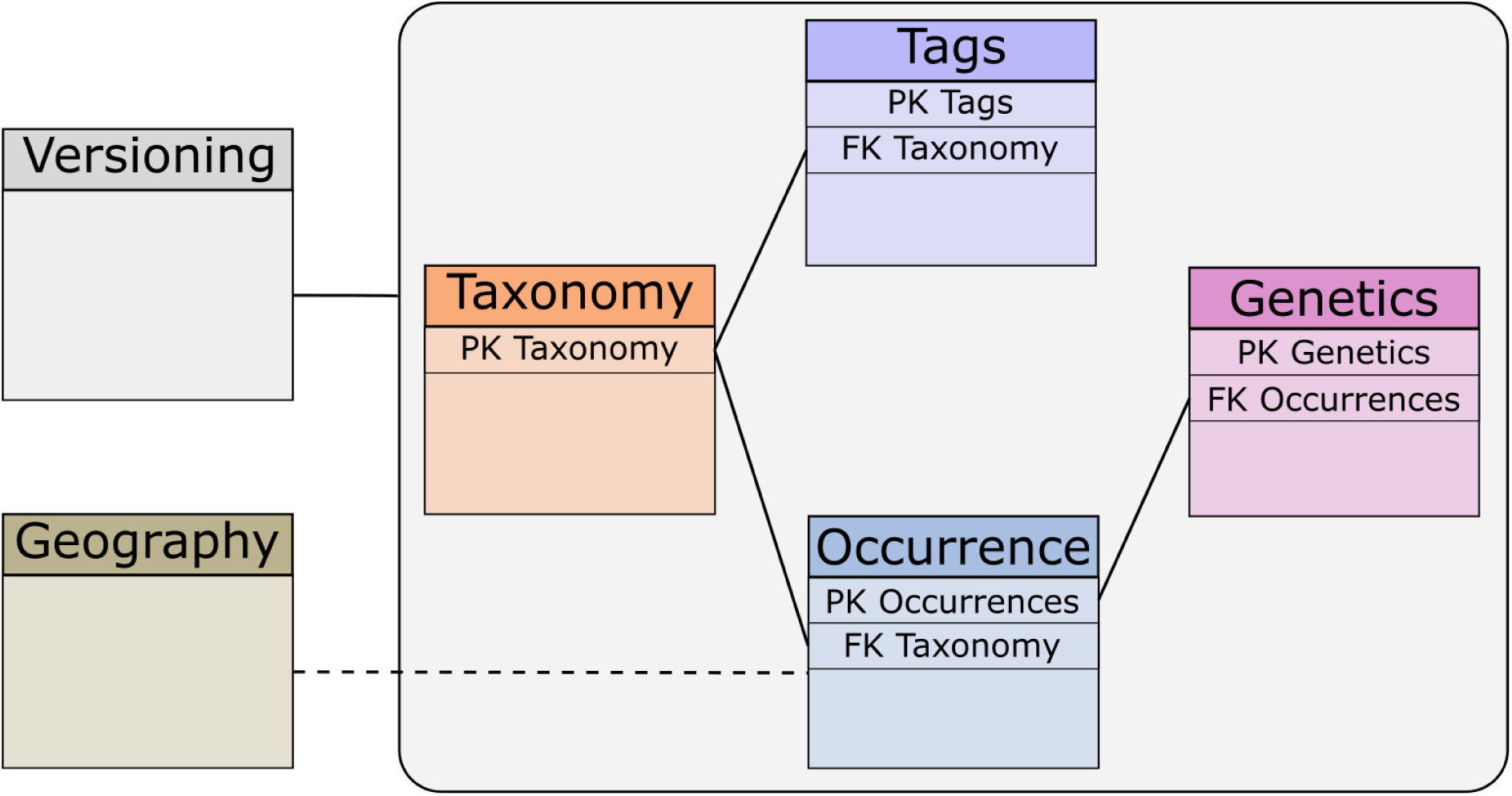
Simplified database schema showing its six thematic modules. Solid lines indicate direct relationships between modules, whereas dashed lines indicate indirect relationships. PK: primary key; FK: foreign key. The full schema is available in the *Supplementary Material S2 (*Figure 1) or can be downloaded from https://osf.io/qwz89/files/utb3m.

The *Taxonomy* module is the centre of our database. This module stores the scientific nomenclature for every taxon included in the database and is organised according with standardised rules of biological nomenclature. We organised the *Taxonomy* module in a hierarchical tree structure where each node represents a taxon that inherits from a common ancestor (the root node; Fig. 3). This parent–child organisation ensures that every taxon is connected to its higher taxonomy, enabling precise and consistent relationships across the entire taxonomic classification. By following this strict hierarchical organisation, we can efficiently navigate and query the taxonomy tree, supporting searches both downward (from higher taxonomic level to lower) and upward (from lower to higher). Additionally, this recursive design removes the need to store the complete higher taxonomic chain for each node. This significantly reduces storage requirements and simplifies maintenance, allowing the system to easily adapt to the dynamic nature of biological classification and its frequent revisions. The taxonomic tree begins with a top ancestor (e.g., Biota, rank Life), followed by eight nested taxonomic levels: kingdom, phylum, class, order, family, genus, species, and subspecies or variety. We chose to include these levels to keep the taxonomy simple and aligned with our objectives. It is important to note that the primary purpose of BiOS is to work as a tool to support biodiversity management, conservation efforts, and the identification of knowledge gaps in biodiversity, rather than for detailed taxonomic purposes. Nevertheless, due to the open-source code, users can add, modify, or remove taxonomic levels in the tree as needed to accommodate specific requirements.

**Figure 3.**
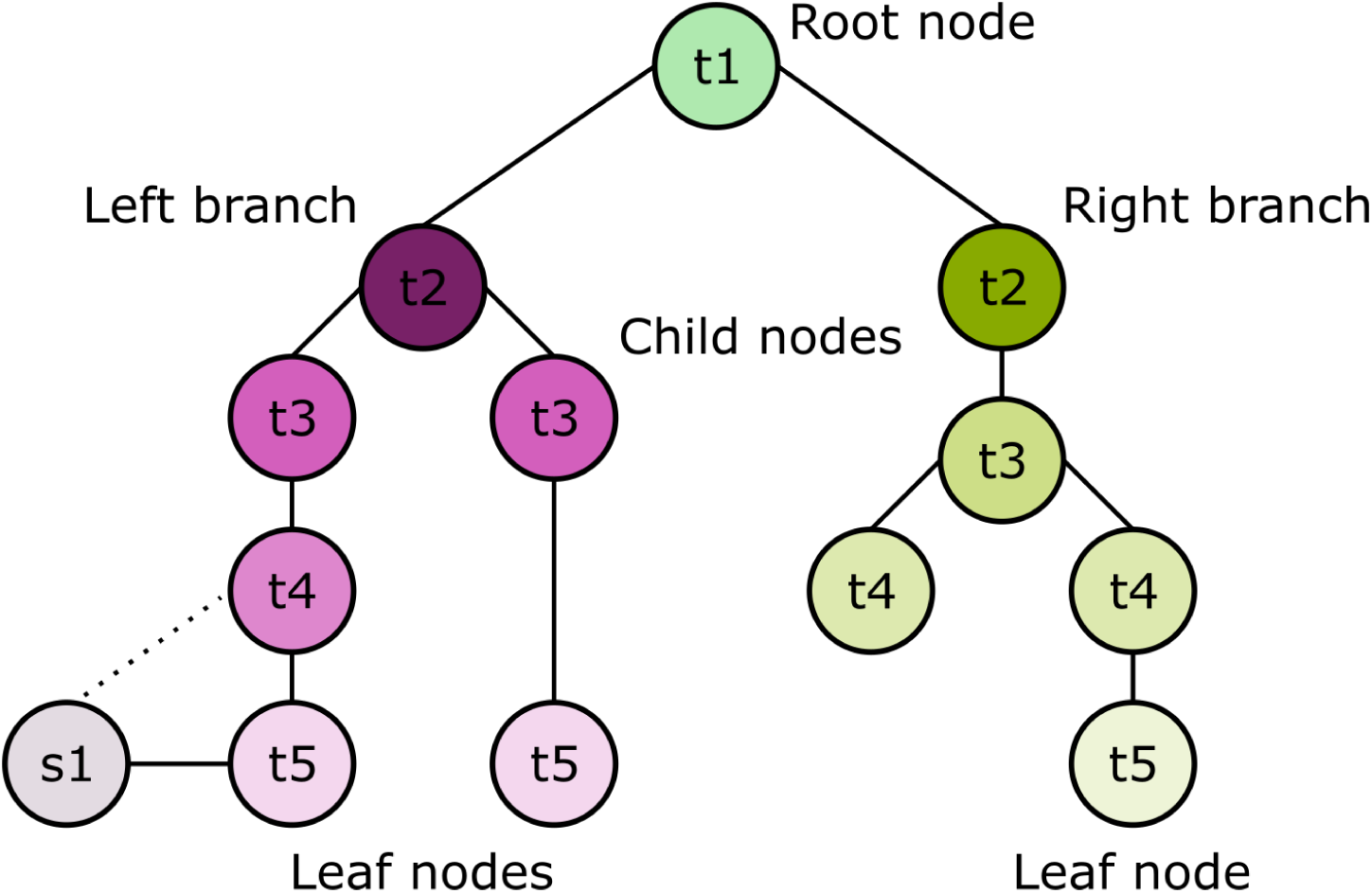
Taxonomic hierarchy scheme. Levels range from “t1”, representing the highest taxonomic rank (the common ancestor or root node), to “t5”, representing the lowest (leaf nodes). “s1” indicates an example of synonym. Circles represent tree nodes. Solid lines indicate the links between nodes. The dashed line represents the connection between the synonym and the parent node.

The *Taxonomy* module also implements features designed to handle taxonomic synonyms. Specifically, a taxonomic node can be assigned the “synonym” attribute and linked to a corresponding “accepted” node. This ensures that all data associated with a synonym are also connected to the branch of the accepted name, allowing access to the metadata connected for both taxon names (Fig. 3, synonym node s1).

Linked to the *Taxonomic* module are the *Occurrence*, *Genetics*, and *Tags* modules, which store complementary metadata associated with a taxon. We implemented a normalised schema to strictly minimise data redundancy by avoiding storing repetitive nomenclature by referencing the *Taxonomy* module through a foreign key. The *Occurrence* module includes information about georeferenced organism records at a specific time. It contains details on where and when a taxon was observed or collected, including geographic coordinates, locality name, dates, and the person or entity that recorded the information. We stored and standardised the geographic coordinates using the WGS84 reference system, ensuring consistency and compatibility with many GIS tools and facilitating data exchange with global biodiversity databases (see *Supplementary Material S*2.2 for more information on the relationships between the fields in the different tables).

The *Genetics* module contains metadata associated with the genetic information of a given taxon occurrence. In the database, we intentionally do not create specific fields to store full genetic sequences to prevent excessive storage demands. Instead, we included fields to store information such as the sequence marker type (e.g., COI, 16S, 18S, etc.), accession identifiers (e.g., GenBank accession number), and other relevant genetic descriptors. In designing this module, we assumed that genetic material is derived from a specimen sampled at a specific location, thus associated with a specific occurrence. Consequently, the *Genetics* module is not directly linked to the *Taxonomic* module, instead, the connection is established indirectly through the *Occurrence* module. We standardised genetic marker names to preserve consistency across the entire database using the same synonym approach as in the *Taxonomy* module.

The *Tags* module was developed to store several types of metadata that complement the ecological and legislative information associated with a taxon. This module supports multiple-region conservation statuses of species (e.g., following the IUCN Red List evaluation such as Least Concern, Near Threatened, etc.), its degree of establishment (e.g., Darwin Core standard), its specific habitats and ecological systems (e.g., as defined by the IUCN Habitats Classification Scheme), as well as its inclusion within national or international directives. Similar to the *Occurrence* module, tags are linked to the *Taxonomy* module.

We prepared the *Geography* module to store geographic geometries, such as spatial polygons provided in shapefile format. This module allows the execution of spatial intersection analyses and supports a variety of geospatial operations. It is also queried by the API to filter occurrence records based on the attributes and spatial relationships of the multilayer polygon features (*Supplementary Material S4.3*).

Finally, we developed the *Versioning* module that implements a dual-layer tracking strategy that facilitates flexible data lifecycle management. The *Batch* model encapsulates a data import event, allowing operations on full datasets, such as the removal of a complete upload history. Whereas, the *Source* model provides granular control over data provenance. This architecture ensures that specific pieces of data or repositories can be isolated and managed independently, without compromising the integrity of the entire batch (*Supplementary Material S3.3*).

### 2.2 API System

We developed an optimised API system to establish a solid communication infrastructure between the database and the web interface, as well as the end users (*Supplementary Material S4*). Following the thematic modules organisation, we implemented six API blocks: Taxonomy, Occurrence, Genetics, Tags, Geography and Versioning. Each block is responsible for managing specific types of data and enabling complex queries (e.g., the Taxonomy endpoint provides access to different taxonomy-related data and performs defined filter operations). To ensure intuitive architecture and rapid data access, API blocks are named to mirror the database’s Thematic modules. Endpoints are organised using a strict, versioned routing structure: */api/{version}/{module}/{function}* (e.g., */api/v1/occurrence/list*). The API framework is built around two main types of endpoints: data retrieval, which handle queries to the database and return specific types of information (e.g., taxonomy of a species), and count, which provide a numerical summary of the requested data (e.g., how many occurrences of a species are stored in the database). The entire framework was developed using Django REST Framework and documented with *drf-yasg* (Fig. 4, drf-yasg, 2023). Documentation on the API endpoints will be available at */api/docs* once the Django environment has been set up (*Supplementary Materials S1*).

**Figure 4.**
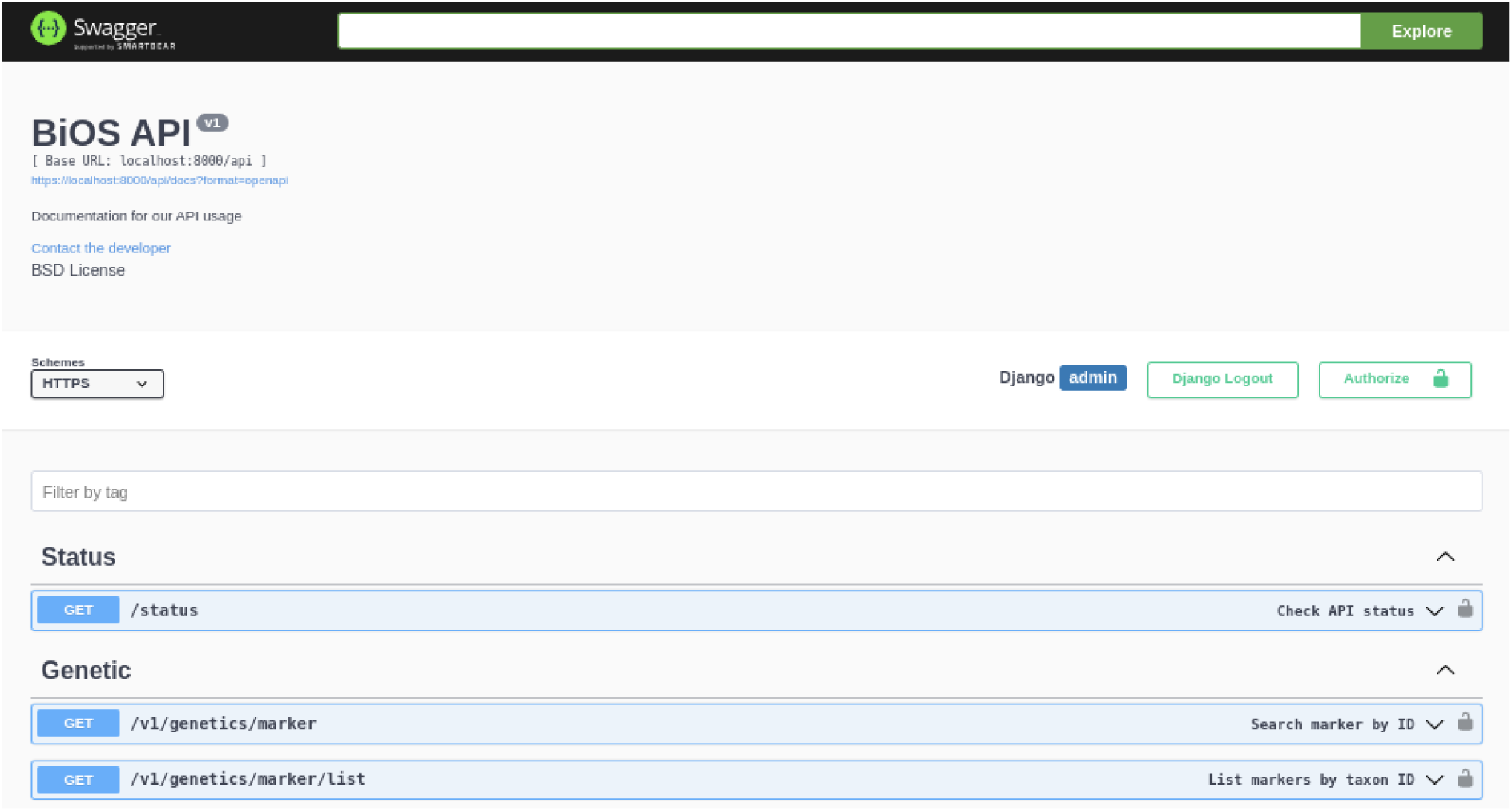
Screenshot of the interactive API documentation interface for the BiOS system, generated using Swagger. This interface exposes the back-end architecture, allowing users to visualise and interact with the API’s resources.

### 2.3 User Interface

To create an intuitive, responsive, and easy-to-use web tool connected with our back-end, we developed the graphical user interface using Next.js. This web development framework provides a powerful solution for building fast and scalable online applications. Among the features that motivated our choice of Next.js (e.g., built-in routing, incremental static regeneration, and strong performance optimisation) is that it notably enables the implementation of server-side–rendered web components. This relatively recent technology improves performance by rendering the initial HTML page on our server rather than in the user’s browser. This feature is particularly useful in our case, and at this stage of the project, especially when handling large amounts of data that could otherwise overload the user’s browser. Additionally, Next.js is a widely adopted framework that benefits from a large developer community, providing extensive third-party packages for web development, and fast support for troubleshooting. For further details on how to deploy the front-end see *Supplementary Material S5*.

This framework also incorporates a powerful caching system. By caching rendered pages and API responses, Next.js reduces the need to repeatedly generate content for frequently accessed resources, minimising server load and accelerating page delivery to users. Therefore, this significantly enhances the performance and responsiveness of our web applications, improving the user experience.

The front-end is organised by adopting a routing strategy based on the */[lang]* directory pattern (eg. */en/about and /es/about* both will render the same content in different languages). This architecture facilitates native internationalisation, ensuring multi-language accessibility across all components through a localised routing system. The platform is structured into four primary dynamic views *Home*, *Taxon*, *Map*, and *Sources* and a static view *About* which provides general information about the platform’s vision and objectives. To navigate between views and change the preferred language, a navigation bar is displayed at the top of the interface and a site map component at the bottom across all pages.

The *Home* view serves as the primary portal for user engagement, featuring a centralised search interface and a summary dashboard that aggregates global database metrics, including total species richness, occurrence counts, and genetic record availability. Upon executing a search for a scientific taxon name, the system redirects to the taxonomic view for the selected taxon. The interface is partitioned into a hierarchical layout, with the lateral panel facilitating nomenclatural navigation through a visual component of the taxonomic tree and associated synonyms, whilst the central header provides critical metadata such as authorship, nomenclatural status, an image of the organism, and unique persistent identifiers. In-depth taxon data view is organised into three thematic modules: the *Overview* tab synthesises ecological attributes including degree of establishment, habitat preferences, and regional conservation status; the *Distribution* tab offers an optimised cartographic distribution of known species occurrences and aggregated data for seasonality, history, and sources of the records; and the *Genetics* tab provides a comprehensive index of related genetic information associated with the taxon.

The *Map* view provides a robust environment for the interactive exploration of species distributions, allowing users to select and visualise multiple taxa simultaneously. Integrated into the front-end, the viewer utilises the “react-map-gl” wrapper with MapLibre (https://maplibre.org/) to efficiently render high-density datasets. The visualisation engine is built on a multi-layer architecture prepared also for 3D environments, supporting the integration of diverse raster sources and utilising raster-dem tiles for dynamic terrain and hillshading (Fig. 5). The component communicates with the RESTful API, querying the specific */occurrences/map* routes, which performs advanced spatial queries on the PostGIS back-end to serve coordinates and metadata accurately. This design ensures that heavy geospatial processing is offloaded directly to the database. Users can interactively query specific records by clicking on individual occurrences to request its metadata and utilise the left-hand panel to filter results based on spatial uncertainty or localities. When locality filters are applied, the database executes spatial intersections between the georeferenced occurrences and the selected geographic polygons defined in the *Geography* module. To support data dissemination, the view includes a utility to export the current geospatial visualisation as a high-resolution image (.png format).

**Figure 5.**
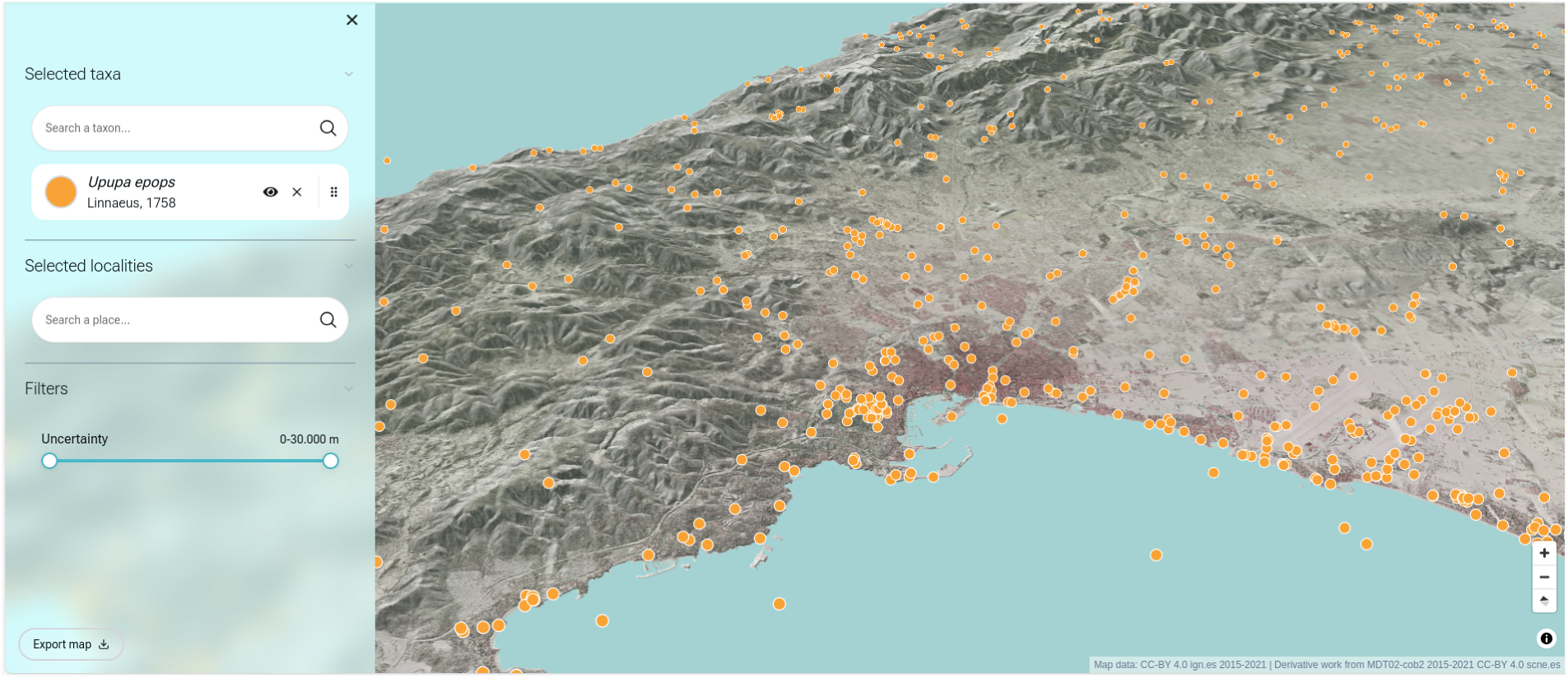
The interactive geospatial visualisation interface view. It allows users to map species occurrences on a 3D terrain model. In this example, the distribution of the Hoopoe (*Upupa epops*) is visualised across the landscape with orange markers. The left-hand panel provides controls for searching specific taxa and localities, as well as a slider to filter records based on spatial uncertainty.

The figures below illustrate a practical example of an instance of BiOS used to create Balearica (Section 3), a regional biodiversity platform (Fig. 5-7).

**Figure 6.**
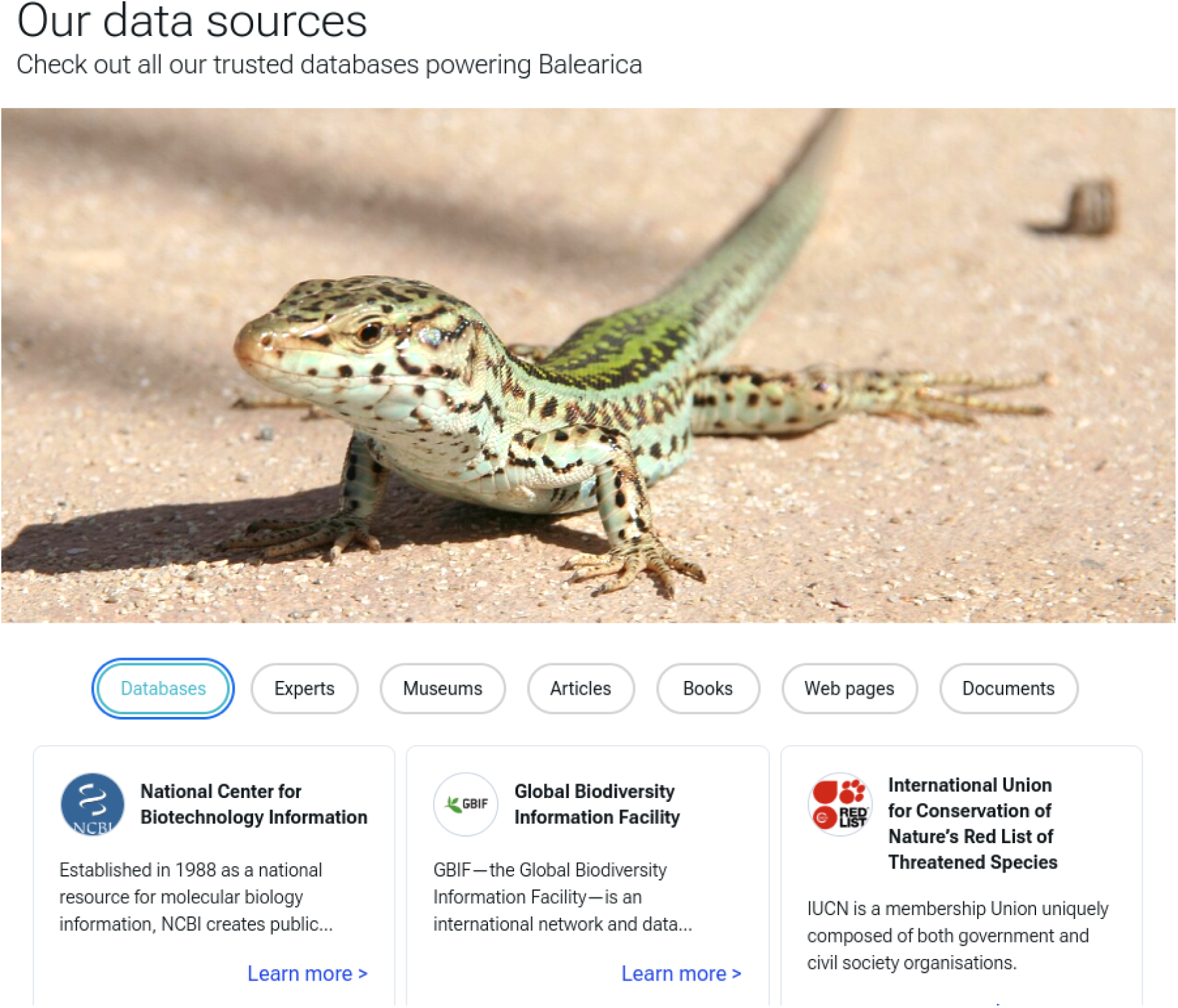
This section lists the external repositories integrated into the portal. As shown in the “Databases” tab, the system aggregates biological records from major global infrastructures. By selecting a repository, the user is redirected to a specific source profile containing the aggregated data and citation information. Lizard photo: Wikipedia; Albert Masats, public domain (PD).

**Figure 7.**
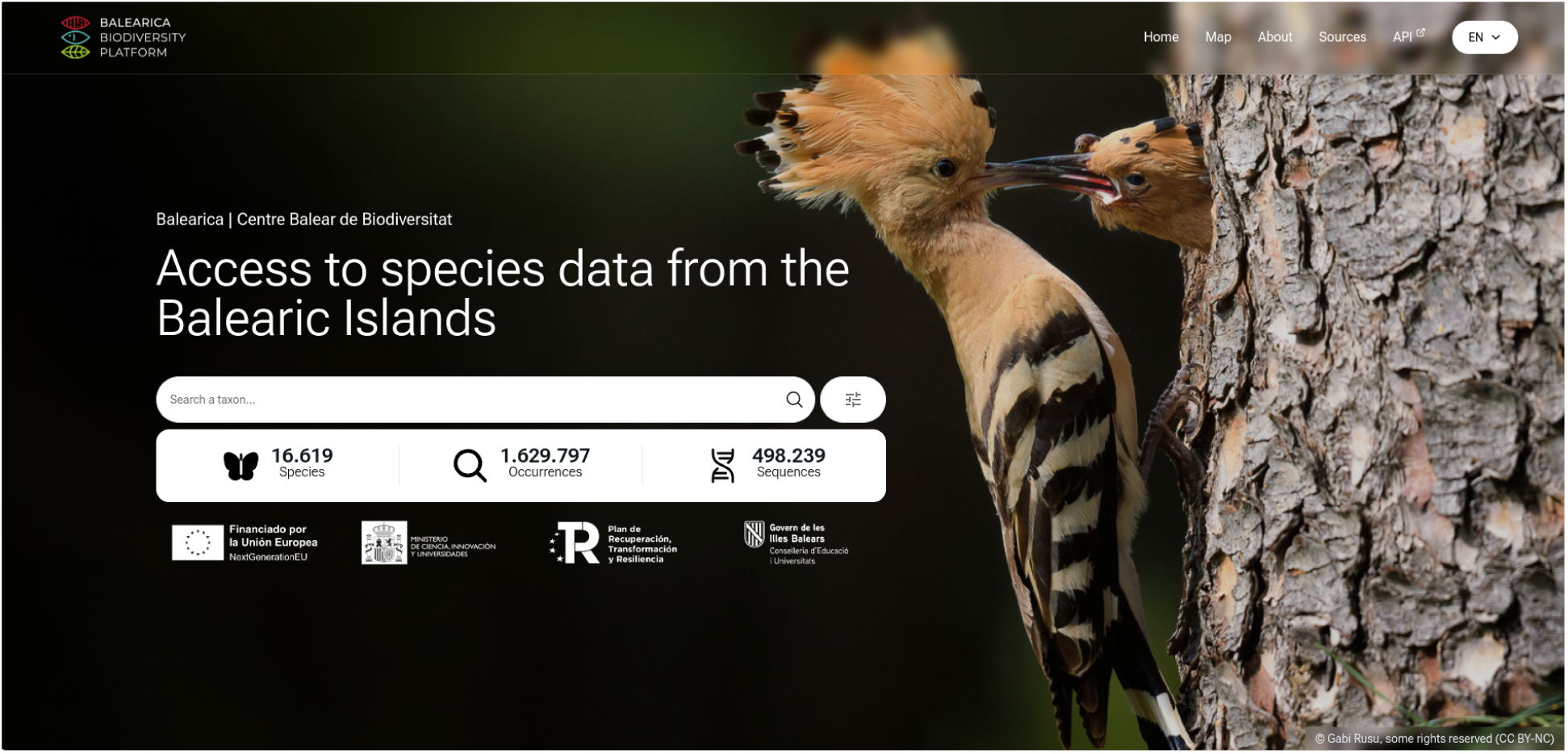
The main landing page of the Balearica web portal, serving as the primary user interface for the BiOS system. Background photo: © Gabi Rusu, some rights reserved (CC BY-NC).

We managed the data transparency through the *Sources* view, which provides a comprehensive registry of all datasets integrated into the platform (Fig. 6). This view is designed to offer an accessible method for identifying and referencing the constituent datasets. To support academic referencing and data validation, each source entry can be expanded to reveal descriptive metadata and summary statistics regarding the specific volume and nature of the records contributed, ensuring that users can accurately identify and cite the primary data providers.

## 3. Example Application: Balearica

To evidence of the system’s capacity, we employed the BiOS framework to develop Balearica, a regional infrastructure that continues to provide information about Balearic biodiversity. Through the integration of heterogeneous data sources, the platform provides a comprehensive and unified view of Balearic biodiversity, supporting scientific research and enabling the study of biological diversity across multiple biological dimensions from taxonomic to genetic. To date (accessed 22 December 2025), Balearica has catalogued more than 16,500 species and integrated approximately 1,600,000 occurrences. In addition, the platform links nearly half a million genetic sequences. Further details on the platform’s architecture, data sources, and methodological approach are available in its official documentation (https://balearica.uib.cat/method/index.html), and the platform can be accessed directly at https://balearica.uib.cat (Fig. 7).

## 4. Discussion

The assessment conducted in Table 1 highlights a persistent structural fragmentation within current biodiversity data infrastructures (Feng et al., 2025; Güntsch et al., 2024). Several existing platforms have evolved as highly specialised systems, each optimised for specific data domains such as taxonomy, species occurrences, genetic information, or conservation status. While this functional specialisation has enabled the development of authoritative repositories within particular fields, it has also resulted in limited cross-domain integration as pointed out by Beyvers et al., 2025. Consequently, researchers and biodiversity managers frequently need to aggregate and reconcile data across multiple independent platforms in order to obtain a comprehensive species profile.

This structural separation poses practical and methodological challenges. Multidisciplinary biodiversity research increasingly requires the simultaneous consideration of taxonomic, ecological, genetic, legislative, and conservation-related information. When these dimensions are distributed across distinct infrastructures, data harmonisation becomes a technically demanding process involving format transformation, taxonomic reconciliation, and repeated API interactions. Such workflows may introduce inconsistencies and reduce analytical reproducibility, particularly in projects operating at regional or local scales with limited technical capacity. With the release of BiOS, we provide a robust, open-source framework that moves beyond the traditional monolithic database model. By strictly adhering to the FAIR principles, BiOS addresses the critical challenge of data heterogeneity identified by Feng et al. (2025). Unlike static repositories, BiOS operates as a dynamic relational engine, harmonising heterogeneous data types such as taxonomic, genomic, and geospatial into a unified framework.

The successful deployment of Balearica (https://balearica.uib.cat) demonstrates the system’s capacity to aggregate and manage high-volume data (over 1.6 million occurrences), validating the framework’s stability and scalability for regional biodiversity observatories. Currently, the landscape of biodiversity informatics is anchored by large-scale infrastructures such as the Atlas of Living Australia (ALA; Belbin et al., 2021) and the Living Atlases community (Lecoq et al., 2017). Whereas the ALA platform is widely considered the “gold standard” for national nodes, its implementation presents a steep technical learning curve. Deploying an ALA stack typically requires significant server infrastructure and a dedicated, highly skilled DevOps team, resources that are often unavailable for many small research centers or regional governments. In contrast, BiOS offers a lightweight solution, delivering robust capability without the heavy infrastructure overhead. Thus, BiOS fills a central niche: allowing institutions to rapidly establish a standards-compliant biodiversity observatory without the massive IT overhead required by global-tier infrastructures.

The design of BiOS is a step forward in addressing the “seven shortfalls” of biodiversity knowledge described by Hortal et al. (2015), particularly the taxonomic (*Linnean*) and distributional (*Wallacean*) shortfalls. The platform’s taxonomic backbone enables experts to identify potential nomenclatural gaps, while advanced geospatial filtering tools may help reveal spatial biases in sampling effort. Furthermore, the integration of a dedicated genetic module enables BiOS to determine which genetic information is available for the taxa stored in the database and to assess whether these data originate from the study region or obtained worldwide. Such integration remains uncommon in regional databases, where genetic data are typically maintained separately from ecological records. This structured framework enables users to identify existing knowledge deficiencies and to support the development of targeted research projects, thereby facilitating the allocation of funding to fill these gaps.

A foundational technical paradigm of the BiOS architecture is the deliberate decoupling of the front-end from the back-end via an exposed API. Traditional biodiversity portals are often tight coupling between data logic and presentation layers, making them difficult for external institutions to adapt or maintain (Mesibov, 2018). By separating these concerns, BiOS offers two main advantages. First, the RESTful API democratises access to the data, allowing researchers to bypass the web interface and interact programmatically with the database for reproducible research and automated meta-analyses. Second, this modularity ensures the longevity of the software; the front-end can be completely redesigned to suit emerging web technologies without disrupting the underlying data integrity.

Beyond academic research, BiOS could serve as a versatile toolbox that bridges the gap between raw scientific data and relevant stakeholders (e.g., public administrations, private environmental agencies, or NGOs), facilitating the acquisition and visualisation of data to enhance and accelerate complex decision-making processes. The inclusion of a legislative labelling module allows the system to cross-reference biological records with local, national, and international protection statutes. This feature is particularly valuable for public administrations, to use the platform as an active management tool, in order to optimise and accelerate decision-making processes. This capability underscores the potential of BiOS to serve not just as a research repository, but as a piece of critical digital infrastructure for governance.

Whilst BiOS offers a comprehensive solution for data management and interoperability, challenges persist regarding data ingestion at scale, particularly concerning the unstructured free text contained in the scrutinised databases (referred to as “latent data” in this article). The results of our platform assessment (Tab. 1), reveal a contrast between biodiversity literature data and actual machine-readability. While taxonomy is generally structured and accessible across all assessed databases, foundational biodiversity data and more complex ecological or biological descriptors frequently suffer from structural bottlenecks. Critical information such as traits, taxonomy metadata, and even georeferenced distributions remain deeply embedded within unstructured text across several major repositories. This reliance on free-text formatting creates a significant barrier to large-scale automated ecological analyses. Consequently, while these databases hold high biological value, leveraging their full scope currently requires substantial secondary extraction efforts, underscoring the need for advanced text-mining solutions or stricter data-entry standardisations within the biodiversity informatics community.

Future development iterations will focus on semi-automating these workflows by integrating biodumpy (Cancellario et al., 2025) to automatically update and retrieve already existing data from large-scale public databases under instance-specific constraints (e.g. the territorial scope). A first step towards mitigating the barrier associated with biodiversity data embedded in scientific and grey literature would be the integration of Specifind (Golomb Durán et al., 2025) to extract species occurrences into a structured machine-readable format. Furthermore, we aim to develop a dedicated module to include biological and ecological traits of species, increasing the multidimensional view of biodiversity. Collectively, these features will form a comprehensive, integrated suite of tools designed to improve biodiversity data management, accelerate analyses, and facilitate a multidimensional understanding of species distributions, ecological traits, and ecosystem dynamics.

Within this context, BiOS does not aim to replace established global infrastructures, nor to replicate the specialised functions of domain-specific repositories. Instead, it represents a shift towards modular, community-driven biodiversity informatics. By providing a free, open-source, and full-stack solution, we aim to lower the technical barrier for institutions worldwide to establish their own biodiversity portals. Whether applied to an archipelago ecosystem like the Balearic Islands or scaled to continental datasets, BiOS offers the structural foundation necessary to turn fragmented records into actionable scientific knowledge.

## Glossary

*Accessible*: It should be clear how metadata and data can be accessed, including any required authentication and authorisation (https://www.go-fair.org/fair-principles).

*Back-end*: server-side part of a software (often a database and RESTful API) that manages the data storage, processing, and application logic.

*Findable*: Metadata and data should be easily accessible to both humans and computers (https://www.go-fair.org/fair-principles/).

*Front-end*: client-facing part of a software that users interact through a graphical user interface such as a web application. It handles the interaction with the back-end via APIs.

*Interoperable*: data need to be integrated with other datasets or metadata, and their format should be compatible with applications or workflows for analysis, storage, and processing (https://www.go-fair.org/fair-principles).

*Interoperability*: ability of distinct databases or systems to exchange information with minimal manual effort and effectively use the information exchanged.

*Linnean shortfall*: the gap between existing species diversity and the species that have been formally described and named. This shortfall also applies to extinct species, which are often underrepresented in the taxonomic record.

*RESTful API*: standardised web interface that allows software components to communicate over HTTP using stateless requests. It enables external systems and applications to access, retrieve, and manipulate data from the back-end in a structured manner, supporting integration with other tools and workflows.

*Reusable*: metadata and data should be accompanied by well-described documentation to enable replication and/or integration in different usages (https://www.go-fair.org/fair-principles).

*Scalability*: ability of a system to handle increasing volumes of data or computational workload.

*Wallacean shortfall*: the lack of knowledge about the geographic distributions of species, even for those that have already been described taxonomically

## Acknowledgements

This study has been partially funded by GOIB/Conselleria d’Educació i Universitats through the project “SINCO2022/18146” and co-funded by the European Union.

This work has been partially funded and promoted by the Comunitat Autònoma de les Illes Balears through the Conselleria d’Educació i Universitats and by the European Union-Next Generation EU/PRTR-C17. I1 (SINCO2022/6717). Nevertheless, the views and opinions expressed are solely those of the author or authors, and do not necessarily reflect those of the Conselleria d’Educació i Universitats, the European Union or the European Commission. Therefore, none of these organisations shall be held liable.

We would like to thank the staff at the Centre Balear de Biodiversitat (CBB-UIB), Laura Bonacho Noguera, and Joan Pons Pons, for their contributions during the development of BiOS. We also thank everyone who has supported the initiative by offering insightful feedback, helpful suggestions, and continued support.

## Authors’ contribution

TC, MC, and EA recognised the need of a modular and multidata biodiversity platform and conceived the conceptual framework to develop BiOS. TC and TGD conceived the back-end architecture, while AR and TGD developed the first version of the back-end. AR and TGD also developed the RESTful API system. TGD developed the front-end, with input and suggestions from all other authors. AJF and TC tested the software. AR, TC, and TGD wrote the original draft and led the manuscript writing. All authors contributed to discussions on the methodology and tested the software. They reviewed and edited the first draft of the manuscript and provided critical input for the final version.

## Conflict of interests

The authors declare no conflicts of interest.

## Notes

https://github.com/centrebalearbiodiversitat/BiOS_backend

https://github.com/centrebalearbiodiversitat/BiOS_frontend

https://osf.io/qwz89/

### Competing Interest Statement

The authors have declared no competing interest.

### Summary of Updates

The main manuscript has been revised, and the supplementary material has been expanded with additional paragraphs.

