## Supplementary Material 1 for "BiOS: An Open-Source Framework for the Integration of Heterogeneous Biodiversity Data"

### Title page

**Article type:** Software and Protocols
**Title:** BiOS: An Open-Source Framework for the Integration of Heterogeneous Biodiversity Data

**Authors:** Alejandro Roldán[1#], Tomás Golomb Durán[1#], Antoni Josep Far[1], Maria Capa[1], Enrique Arboleda[1], Tommaso Cancellario[1]*

[#] Equally shared first authorship

**Affiliation:**
[1]Centre Balear de Biodiversitat, Universitat de les Illes Balears, Palma, Spain

*** Corresponding author:** Tommaso Cancellario
 -
CBB - University of the Balearic Islands (Spain)

ORCID:
AR 0009-0009-5040-1009
TGD 0009-0003-0430-2073
JAF 0009-0008-8869-6482
MC 0000-0002-5063-7961
EA 0000-0001-9800-1674
TC 0000-0002-6637-4764

### Supplementary Materials

Here we aim to provide the technical documentation to deploy the BiOS framework on own or a third-party server.

The supplementary material is divided into the following main sections.

- S1: Installation and Deployment Guide
- S2: Detailed operation of the database
- S3: Data Loading and Bulk Ingestion Workflow
- S4: Technical API Documentation
- S5: Front-end deployment

### S1. Back-end deployment

To ensure consistency between development and production environments, we used Docker (Docker Inc., 2025) containers to deploy the BiOS framework. This approach creates an isolated system for both environments, ensuring an efficient management of dependencies [i.e., PostGIS (PostGIS Developers, 2023) and Django (Django Software Foundation, 2023)] and increasing the reproducibility of the product.

#### S1.1. Environment Dependencies

The BiOS framework is based on the following Python (Van Rossum and Drake, 2009) packages (Tab. 1), all defined in the *requirements.txt* file.

**Table 1.** Required Python packages.

| Dependency | Version | Primary Function | Reference |
|---|---|---|---|
| Django | 4.2.5 | Main back-end framework | https://www.djangoproject.com. |
| django-mptt | 0.16.0 | Management of hierarchical structures (Taxonomy, Geography) | https://github.com/django-mptt/django-mptt |
| djangorestframework | 3.15.1 | Creation of the RESTful API (Christie, 2023) | https://www.django-rest-framework.org/ |
| djangorestframework-gis | 1.1 | Serialisation of PostGIS geospatial data | https://github.com/openwisp/django-rest-framework-gis/tree/master |

| | | | |
|---|---|---|---|
| drf-yasg | 1.21.7 | Automatic generation of API documentation (Swagger) | https://github.com/axnsan12/drf-yasg |
| psycopg2-binary | 2.9.5 | Connection of Django with PostgreSQL (The PostgreSQL Global Development Group 2023) | https://pypi.org/project/psycopg2-binary/ |
| geopandas | 1.0.1 | Python module for advanced handling and analysis of spatial data (Jordahl et al., 2020) | https://geopandas.org/ |

### S1.2. Quick Installation Steps

Deploying the back-end service locally (i.e., the database and API) requires the installation of Docker (to create and run containers for each component) and Docker Compose (to start and manage all containers together using a single configuration file).

**Note:** The code blocks shown below are valid for Unix-based operating systems. If you are using a different operating system, you will need to use the corresponding commands.

The procedure for launching the Docker container and deploying the Django application is outlined below.

1. Open a terminal and clone the repository from GitHub:

```
git clone
https://github.com/centrebalearbiodiversitat/BiOS_backend.git
cd BiOS_backend
```

2. Configure the environmental variables to store common settings across the different environment setups (development and production). To configure these variables, rename the file located in the directory *BiOS/* from *.env_template* to *.env*. Then, modify the parameter of your *.env* file following the indications provided in Table 2.

**Table 2.** Parameters and their detailed descriptions for compiling the *.env* file.

| Field | Values | Description |
|---|---|---|
| DEBUG | true/false | Controls whether the application runs in debug mode (true). Detailed errors are shown and files are automatically reloaded when saved. Must be false in production. |

| | | |
|---|---|---|
| DJANGO_SECRET_KEY | Random alphanumeric string | Secret key used by Django for security. It is critical. It must be unique and confidential. |
| SITE_URL | URL or hostname (e.g., localhost, 127.0.0.1, mydomain.com) | The base URL or domain where the application is hosted. Automatically populates needed Django configuration (ALLOWED_HOSTS, SITE_URL, and CORS). |
| DB_HOST | Hostname or IP address of the database server (e.g., postgres, 127.0.0.1) | The address or name of the container/service where the database is hosted. Set by default to the docker postgis service. |
| DB_PORT | Port number (e.g., 5432 for PostgreSQL) | The network port on which the database server is listening for connections. Must match the docker-compose file. |
| POSTGRES_DB | Database name (e.g., bios_db) | Desired name of the specific database within the server to which Django will connect. |
| POSTGRES_USER | Text string | Desired username with permissions to access the database. |
| POSTGRES_PASSWORD | Text string | Desired password corresponding to the database user. |

3. Download the content from the following OSF link: https://osf.io/qwz89/files/osfstorage

4. Next, replace the 'cbb' folder with the 'cbb' folder found in the downloaded OSF content.

5. Copy and paste the 'docker-compose.local.yml' file initially located into the root of the *BiOS/* project.

   Once the .yml file is stored in the BiOS/ folder, you can build the Docker image (including all dependencies) and launch the services defined in the production (*docker-compose.yml*) or development environment (*docker-compose.local.yml*) running the commands below.

```
# Development
sudo docker compose -f docker-compose.yml -f
docker-compose.local.yml up --build
```

```
# Production
sudo docker compose -f docker-compose.yml up --build
```

The difference between the two commands, lies in the *-f* parameter. While the production environment uses only the basic configuration, the development environment includes additional settings that enhance the developer experience.

6. Open a second terminal. Now we need to open a bash session inside the Docker container to run the command-line for administrative tasks.

```
# Open a bash session inside the Django container (exec terminal)
sudo docker compose -f docker-compose.local.yml exec django bash
```

7. Once the Django service is running, we can execute the Python commands *makemigrations* and *migrate*.

   The first command is useful to compare the current state of the database defined by the *models.py* files contained in each Django /*app* (see S2.1 for further details) with the previous configuration. Whereas, the second is useful to apply possible changes and update the database schema. The commands *makemigrations* and *migrate* are necessary when setting up the application for the first time, since the database structure and table relationships must be defined. However, they are also useful whenever new fields will be added, tables will be deleted, or relations will be modified.

```
# Create migration files for any model changes
python manage.py makemigrations

# Apply the migrations to update the database schema
python manage.py migrate
```

8. Finally, to interact with the database, it is necessary to create a superuser. The superuser is needed because it provides full administrative access to the Django admin site application. Allowing to manage all data models, users, and permissions through the Django admin web user interface.

```
python manage.py createsuperuser
```

API back-end and Swagger/OpenAPI documentation are available at *http://localhost:8000/api/v1/{app}* and *http://localhost:8000/api/docs/*, respectively. While Django admin site at *http://localhost:8000/admin*

### S2. Detailed operation of the database

To develop the BiOS database architecture, we took advantage of Django's Object-Relational Mapping system (ORM), which allows us to define the structure of the database using Python classes (referred to as Models in Django) instead of writing raw SQL code. This high level of coding abstraction facilitates data portability and security. We have organised the project into six modular ‘applications’ (hereafter app), each of which encapsulates a specific thematic domain and follows a distinct computational logic. Specifically, the applications are: taxonomy, occurrences, genetics, tags, geography, versioning. These applications interact with each other through Primary Keys (PK) and Foreign Keys (FK) relationships. Broadly here we define a PK as a unique identifier for each record in a table, while a FK as a field connection field useful to establish a link between two applications.

Taxonomic information (taxonomy app) serves as the central node to which observations (occurrences app), genetic data (genetics app), and other complementary attributes (tags app) are linked. The geography thematic module (geography app) is connected with the occurrence app, and it provides the spatial infrastructure necessary to organise the territory hierarchically and validate the location of records. We also included a cross-cutting versioning thematic module (versioning app) to ensure the traceability of each record stored in the database.

Finally, we based the system architecture on the use of abstract models that provide standardised functionality to the rest of the tables. These abstract tables avoid redundancy and are designed to store well-defined data, which facilitates overall database management and updates. All interactions between tables and their fields can be seen in the following diagram (Fig. 1).

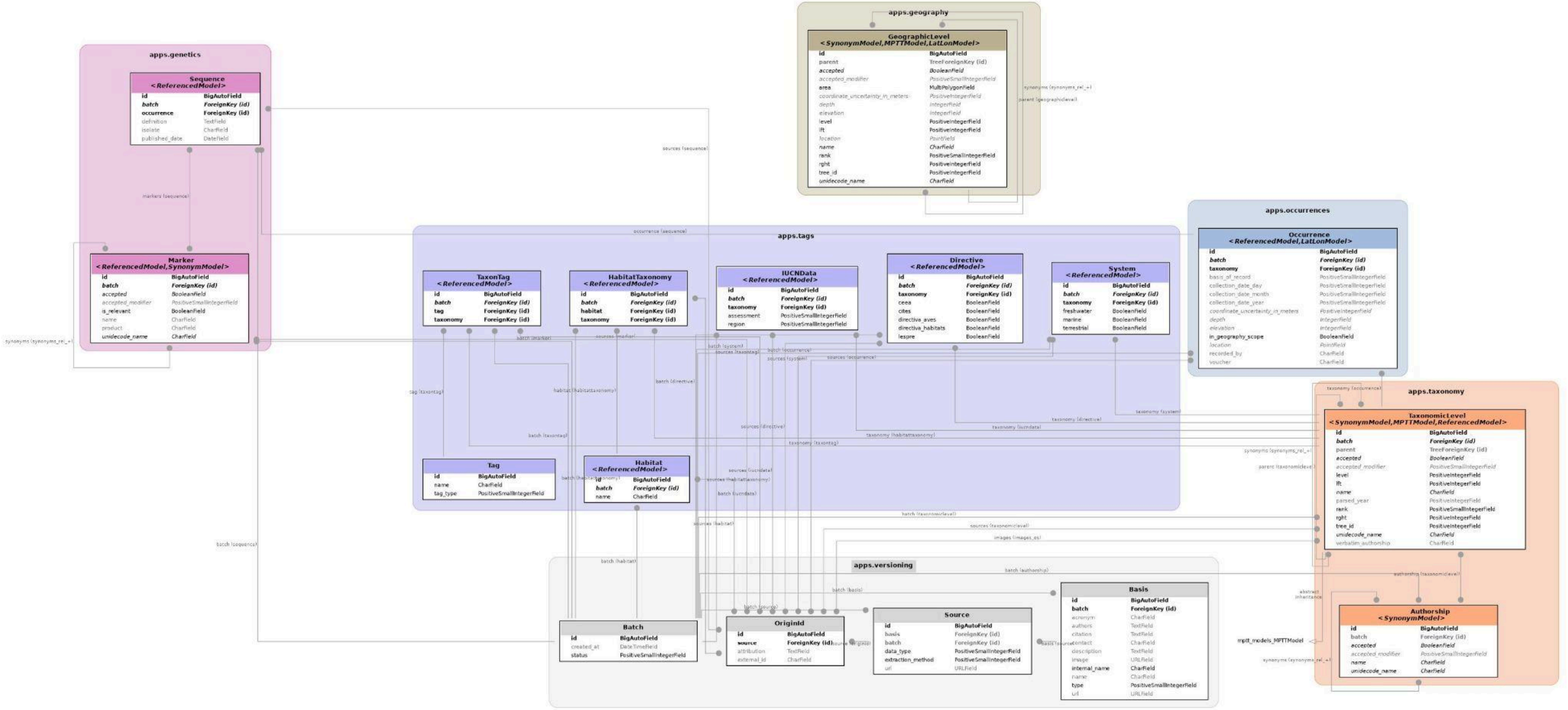


**Figure 1.** Relational architecture of the system using color clusters that delimit its functional thematic modules. Each thematic module is detailed in rectangular nodes that specify its name, abstract inheritance (between < >), and technical attributes. Dependencies are represented by continuous edges where a solid dot identifies the source table of the foreign key (multiple cardinality end) to the referenced entity, while hollow triangular-tipped arrows denote inheritance from abstract base classes. The labeling on the lines reflects the attributes of the source code, and the polyline routing avoids visual overlap, highlighting the versioning thematic module (grey) as the horizontal traceability axis around which the metadata (tags, purple), geography (beige), genetics (pink), occurrences (blue), and taxonomy (orange) thematic modules are articulated. High resolution image available at: https://osf.io/qwz89/files/utb3m.

### S2.1. Details of the database tables

Below, we provide the technical definitions of the fields used in the applications, including the field name, data type (following Django ORM conventions; see Tab. 3), and a brief technical description of each. Additionally, when applicable, for each table we indicate in parentheses (just below the table name) the parent models it inherits from. Only fields unique to each model are listed in the tables below.

**Table 3.** Table of abbreviations for Django ORM data types.

| Acronym | Description |
|---|---|
| FK | ForeignKey |
| MTMF | ManyToManyField |
| TFK | TreeForeignKey |
| CF | CharField |
| TF | TextField |
| IF | IntegerField |
| PIF | PositiveIntegerField |
| PSIF | PositiveSmallIntegerField |
| BF | BooleanField |
| DTF | DateTimeField |
| PF | PointField |
| MPF | MultiPolygonField |
| URLF | URLField |

##### Base abstract models

We implemented multiple base models (Tab. 4) using Django's abstract feature to ensure and reuse multiple capabilities across the end-models:

- Data integrity: Enforced by the *ReferencedModel*, which mandates that every piece of entered data is linked to a quality control batch and its original external sources to guarantee full traceability.

- Handle nomenclatural ambiguity: The *SynonymModel* provides the logic to manage nomenclatural ambiguity in fields such as a taxon or genetic marker name, allowing any record to be classified as either ‘accepted’ or ‘synonymous’ and linked to the other representants.
- Spatial management: Unified through the *LatLonModel*, this model equips any entity with native geospatial capabilities via PostGIS, enabling the secure storage of precise coordinates alongside their uncertainty, elevation, and depth metadata.

**Table 4.** Description of the fields in the tables belonging to the base abstract models.

| Table | Field | Data type | Description |
| --- | --- | --- | --- |
| LatLonModel | location | PF | PostGIS geospatial field that stores the geographical location (WGS84 coordinates). |
| | coordinate_uncertainty_in_metres | PIF | Uncertainty radius of the coordinate in metres. |
| | elevation | IF | Elevation or altitude above sea level. |
| | depth | IF | Depth below water level or surface. |
| ReferencedModel | batch | FK | Link to the control batch. |
| | sources | MTMF | External sources of origin for the record (allows multiple external IDs). |
| SynonymModel | accepted | BF | Indicates whether this entity is the accepted record or name (True) or a synonym (False). |
| | accepted_modifier | PSIF | Status modifier (e.g. PROVISIONAL, AMBIGUOUS). |
| | synonyms | MTMF | List of other entities or names considered synonyms. |

##### Thematic modules

The Taxonomy thematic module is the backbone of the entire database, with all other modules linked to it. We modelled the taxonomic tree in the *TaxonomicLevel* table, where we used a hierarchical algorithm (Modified Preorder Tree Traversal - MPTT) implementing django-mptt (django-mptt, 2024) that allows us to navigate efficiently through its structure (from life to variety). To ensure data standardisation and bibliographic consistency, scientific name authorities are stored in a separate authoring table (*Authorship*) linked to the taxon by external keys. This relational design centralises citation data, eliminating redundancy and avoiding inconsistencies arising from duplicate text strings (Tab. 5).

**Table 5.** Description of the fields in the tables belonging to the Taxonomy thematic module.

| **Table name** | **Field name** | **Data type** | **Field description** |
|---|---|---|---|
| TaxonomicLevel (SynonymModel, MPTTModel) | rank | PSIF | Taxonomic rank (e.g., GENUS=5). |
| | verbatim_authorship | CF | Normalised taxon's authorship. |
| | parsed_year | PSIF | Year of publication of the authorship, extracted and standardised. |
| | authorship | MTMF | Link to the authorship record in the Authorship table. |
| | parent | TFK | Hierarchical link to the superior level (MPTT implementation). |
| | images | MTMF | List of associated images. |
| Authorship (SynonymModel) | batch | FK | Link to the control batch. |

The Occurrences thematic module serves as a bridge linking each observation to the corresponding taxonomic name stored in the table *TaxonomicLevel*. The *Occurrence* table (Tab. 6) by inheriting the spatial capabilities of the *LatLonModel*, can store the exact geographical position of a specific occurrence and validate whether it falls within our study area. It also supports detailing metadata such as the origin of the record (e.g., herbarium specimen, human observation, citation, etc.), the person or entity that collected the specimen, and the collection date.

**Table 6.** Description of the fields in the tables belonging to the Occurrences thematic module.

| **Table name** | **Field name** | **Data type** | **Description** |
|---|---|---|---|
| Occurrence (ReferencedModel, LatLonModel) | taxonomy | FK | Link to the TaxonomicLevel table. |
| | voucher | CF | Unique identifier of the physical sample (e.g., herbarium or museum number). |
| | recorded_by | CF | Name of the person or entity who made the observation/collection. |
| | collection_date_year | PSIF | Year of occurrence record. |

| | | | |
|---|---|---|---|
| | collection_date_month | PSIF | Month of occurrence record. |
| | collection_date_day | PSIF | Day of occurrence record. |
| | basis_of_record | PSIF | Record type (e.g., HUMAN_OBSERVATION = 4). |
| | in_geography_scope | BF | Indicates whether the observation falls within the defined geographical areas. |

The Genetics thematic module contains tables dedicated to storing genetic metadata. It is structured into two tables: *Sequence* and *Marker* (Tab. 7). The *Sequence* table is related to the *Occurrence* table through the *occurrence* field, which stores the ID of the occurrence from which the sample was extracted for sequencing. Moreover, in this table, we stored metadata, such as the different markers sequenced from that sample (*markers*) and the date of publication of the sequence (*published_date*).

To standardise genetic marker names and ensure the use of a controlled vocabulary, we use the *Marker* table. This model inherits from *SynonymModel* that allows us to create a catalog of standardised genetic markers (e.g., accepted name: COI; synonyms: CO I, COI-5P, COX1, COXI, CO1, etc.), thereby providing more robust and reliable responses when querying the database.

The relationship between genetic information and marker names is flexible, allowing us to associate to a genetic record multiple markers.

**Table 7.** Description of the fields in the tables belonging to the Genetics thematic module.

| Table | Field | Data type | Description |
|---|---|---|---|
| Sequence (ReferencedModel) | occurrence | FK | Link to the Occurrence record from which the sequenced sample was extracted. |
| | isolate | CF | Sequence isolation code. |
| | definition | TF | Sequence description. |
| | published_date | DF | Sequence publication date. |
| | markers | MTMF | Link to the associated genetic markers. |
| Marker (ReferencedModel, SynonymModel) | name | CF | Marker name (e.g., COI). |
| | product | CF | Marker resulting product. |

The Tags thematic module includes seven tables and is designed to store a wide range of information that enriches taxon metadata (Tab. 8). We developed this module to manage diverse types of information, ranging from ecological details (e.g., habitat) to national and international legislation.

**Table 8.** Description of the fields in the tables belonging to the Tags thematic module.

| Table | Field | Data type | Description |
|---|---|---|---|
| System (ReferencedModel) | taxonomy | FK | Link to the TaxonomicLevel table. |
| | freshwater | BF | Presence in freshwater. |
| | marine | BF | Presence in salt water. |
| | terrestrial | BF | Presence on land. |
| Tag | name | CF | Name of the tag (e.g., "Established"). |
| | tag_type | PSIF | Tag type (e.g., degreeOfEstablishment=0). |
| TaxonTag (ReferencedModel) | taxonomy | FK | Link to the TaxonomicLevel table. |
| | tag | FK | Link to the Tag table. |
| Habitat | name | CF | Name of the standardised habitat type. |
| IUCNData (ReferencedModel) | taxonomy | FK | Link to the TaxonomicLevel table. |
| | assessment | PSIF | Conservation status (EN, VU, CR, etc.). |
| | region | PSIF | Geographical region for the assessment. |
| HabitatTaxonomy (ReferencedModel) | taxonomy | FK | Link to the TaxonomicLevel table. |
| | habitat | FK | Link to the Habitat table. |
| Directive | taxonomy | FK | Link to the TaxonomicLevel table. |
| | cites | BF | Indicator in the Convention on International Trade in Endangered Species of Wild Fauna and Flora. |
| | ceea | BF | Inclusion in the Spanish Catalogue of Threatened Species. |
| | lespre | BF | Inclusion in the Spanish List of Wild Species under Special Protection Regime. |
| | directiva_aves | BF | Inclusion in the European Union's Birds Directive. |

| | | | |
|---|---|---|---|
| | directiva_habitats | BF | Inclusion in the European Union's Habitats Directive. |

To manage geospatial data, we created the Geography thematic module, which is composed of a single table. Spatial infrastructure is managed using the *GeographicLevel* table (Tab. 9). As with taxonomy, we implement MPTT hierarchical logic here to model administrative and physical divisions of the territory. We store complex geometries (polygons) in the *area* field, allowing us to nest these areas within each other. This design facilitates powerful spatial queries, allowing us to filter all occurrences that fall within a specific municipality or automatically validate input coordinates against official boundaries.

**Table 9.** Description of the fields in the tables belonging to the Geography thematic module.

| Table | Field | Data type | Description |
|---|---|---|---|
| GeographicLevel (SynonymModel, MPTTModel, LatLonModel) | rank | PSIF | Geographical rank (e.g., ISLAND, MUNICIPALITY). |
| | parent | TFK | Hierarchical link to the superior geographical level. |
| | area | MPF | Area geometry (PostGIS). |

To manage data traceability, we designed the Versioning thematic module which contains four tables (Tab. 10). The *Basis* table is used to store general data source information (such as the name or type of data source). This information is specified in the *Source* table, which records the specific sources. Granularity is achieved with the *OriginId* table, which stores the original external identifier for each entry. Finally, we group all this flow in the *Batch* table, which manages the loading of data packages in batches with approval statuses (pending, accepted, or rejected), giving us total control over the lifecycle and quality of the information entering the system.

**Table 10.** Description of the fields in the tables belonging to the Versioning thematic module.

| Table | Field | Data type | Description |
|---|---|---|---|
| Batch | created_at | DTF | Date of data batch creation. |
| | status | PSIF | Status of the batch (PENDING, ACCEPTED, REJECTED). |
| Basis | internal_name | CF | Internal key name for fast queries (e.g., "GBIF", "COL"). |
| | name | CF | Full name of the source. |

| | | | |
|---|---|---|---|
| | acronym | CF | Acronym of the source (e.g., "GBIF"). |
| | type | PSIF | Classification of the source type (e.g., JOURNAL_ARTICLE). |
| | url | URLF | URL of the source's main page. |
| | image | URLF | Path to the source's image or logo. |
| | description | TF | Detailed description of the source. |
| | authors | TF | Main authors or collaborators of the source. |
| | citation | TF | Citation text for the source. |
| | contact | CF | Contact information for the source. |
| | batch | FK | Link to the control batch to which the source belongs. |
| Source | extraction_method | PSIF | Method of data extraction from the source. |
| | data_type | PSIF | Type of data contained in the source (e.g., TAXON, SEQUENCE, etc). |
| | url | URLF | Specific URL where the data from the source was consulted or downloaded. |
| | batch | FK | Link to the control batch to which the source belongs. |
| | basis | FK | Link to the Basis table. |
| OriginId | external_id | CF | External ID of the record (e.g., GBIF ID). |
| | source | FK | Link to the Source table. |
| | attribution | CF | Specific data attribution or citation text. |

### S2.2 Definition of Database Relationships

The BiOS framework uses Foreign Key relationships to ensure that associated data (e.g., an occurrence) maintain consistency with reference data (e.g., taxonomy). Here, we detailed the relationship that links the PK and the FK across the Django Applications (Tab. 11).

**Table 11.** Details the Foreign Key and Many to Many fields relationships implemented in the framework.

| **Source Table** | **Field** | **Type** | **Target Table** | **Implementation Notes** |
|---|---|---|---|---|
| Authorship | batch | FK | Batch | Links the authors to a control batch. |
| TaxonomicLevel | parent | TFK | TaxonomicLevel | Association with its superior geographical level. |

| | | | | |
|---|---|---|---|---|
| | authorship | MTMF | Authorship | Links taxonomy with its author. |
| | images | MTMF | OriginId | Links taxonomy with the original ID of the associated image. |
| Occurrence | taxonomy | FK | TaxonomicLevel | Links the observation to the target species. |
| Sequence Sequence | occurrence | FK | Occurrence | Links the sequence to the occurrence specimen. |
| | markers | MTMF | Marker | A sequence can have multiple markers and vice-versa. |
| System | taxonomy | FK | TaxonomicLevel | Associates the system with the species. |
| TaxonTag | taxonomy | FK | TaxonomicLevel | Links a type of tag to the target species. |
| | tag | FK | TaxonomicLevel | Connect an instance of tag with its type. |
| IUCNData | taxonomy | FK | TaxonomicLevel | Associates the conservation status with the species. |
| HabitatTaxonomy | taxonomy | FK | TaxonomicLevel | Links the type of habitat to the target species. |
| | habitat | FK | Habitat | Connect an instance of habitat with its type. |
| Directive | taxonomy | FK | TaxonomicLevel | Links the directive to the target species. |
| GeographicLevel | parent | TFK | GeographicLevel | Association with its superior geographical level. |
| Basis | batch | FK | Batch | Links the basis to a control batch. |
| Source | basis | FK | Basis | Links the source (e.g., "Smith 2020") to the source type (e.g., "Journal Article"). |
| | batch | FK | Batch | Links the source to a control batch. |
| OriginId | source | FK | Source | Links the external ID to the bibliographic or institutional source. |

### S3. Data Loading and Bulk Ingestion Workflow

To load a bulk of data you can run the *load_db.sh* script in a bash terminal inside Django's Docker container from the command line (see S1.2). If the data are properly stored and organised (follow *S1.2*), the database will be automatically populated. Essentially, the *load_db.sh* script invokes two types of commands, named *load_*.py* or *populate_*.py*, which perform multiple validation steps, resolve foreign keys, and ensure automatic traceability throughout the ingestion process. The difference between them is that while 'load_*.py' loads data from third-party files, 'populate_*.py' inserts data that is predefined in the command code itself. However, to increase the flexibility for the data loading, we also provide the option to populate or modify specific tables using the *load_*.py* and *populate_*.py* scripts, as described in Table 12 - 15.

```
sh load_db.sh
```

#### S3.1. Structured Loading Workflow

The *load_db.sh* script establishes the required execution order for bulk data loading. Data will be loaded in three main phases, ensuring that the base data in Geography and Taxonomy thematic modules, exists before attempting to associate specific taxon records.

> **Note**: To facilitate understanding of the process, a sample data file is attached for each script. These files reflect the structure used for the practical case of the paper, whose data is referenced to the Balearic Islands (Spain). Therefore, all geographical and taxonomic entities shown in the examples will refer to this scope.

**Phase I: Load data in the Geography thematic module**

This phase focuses on populating the *GeographicLevel* table with spatial geometries organised with a top-down territorial hierarchy (Tab. 12). This phase is critical because the Occurrence table will use these geometries for spatial filtering. The *load_gadm.py* script is invoked sequentially to load spatial units with a nested hierarchy (e.g., from Autonomous Community, to Municipalities).

```
# Geography
echo "Loading Autonomous communities..."
python manage.py load_gadm data/GIS/IDEIB_AC/*uncertainess*/*.shp
echo "Loading Islands..."
python manage.py load_gadm data/GIS/IDEIB_islands/*uncertainess*/*.shp
echo "Loading Municipalities..."
python manage.py load_gadm data/GIS/IDEIB_municipalities/*uncertainess*/*.shp
echo "Loading Populations..."
python manage.py load_gadm data/GIS/CNIG_poblaciones/*uncertainess*/*.shp
```

**Table 12.** Details of the commands, files, and paths where the files must be stored for uploading the geographical information, using the *load_db.sh* script.

| I. Geographical Infrastructure (PostGIS) | | | | |
|---|---|---|---|---|
| **Data loaded** | **Command (Django)** | **File path (from load_db.sh)** | **Format File** | **Content Purpose** |
| Autonomous Communities | load_gadm | data/GIS/IDEIB_AC/*uncertainess*/*.shp | .shp | Loads the geometries and names of the geographical units (GeographicLevel). |
| Islands | load_gadm | data/GIS/IDEIB_islands/*uncertainess*/*.shp | .shp | |
| Municipalities | load_gadm | data/GIS/IDEIB_municipalities/*uncertainess*/*.shp | .shp | |
| Settlements | load_gadm | data/GIS/CNIG_poblaciones/*uncertainess*/*.shp | .shp | |

##### Phase II: Base Catalogues and Tags

This phase initialises the reference catalogues that will be used by the traceability system (Versioning) and the data attribution thematic module (Tags). Therefore, reference tables such as *Basis*, *Tag*, and *Habitat* will have the reference value that you can later associate in Phase III with other data such as the name of the database from which an occurrence originates, or the habitat of a species (Tab. 13).

The *load_basis.py* script is used to load general information of the sources as the name (e.g., "GBIF", "NCBI", "COL" or "OBIS"). While *populate_tags.py* and *populate_habitats.py* scripts are executed without input parameters, as they are programmed to initialise the system with basic controlled vocabularies which will then be linked to their corresponding species. In this way, an instance of each type of tag or habitat is created, which will subsequently be connected to each of the species associated with it (this connection occurs in Phase III-A).

The correct execution of this phase is indispensable, as the origin of an occurrence cannot be traced if its source (*Basis*) does not exist.

```
#Populate
echo "Populating basis..."
python manage.py load_basis data/basis.json
echo "Populating tags..."
python manage.py populate_tags
```

```
echo "Populating habitats..."
python manage.py populate_habitats
```

**Table 13.** Details of the commands, files, and paths where the files must be stored for uploading using the *load_basis, populate_tags and populate_habitats* commands.

| II. Base Catalogues and Tags | | | | |
|---|---|---|---|---|
| **Data loaded** | **Command (Django)** | **File path (from load_db.sh)** | **Sample File** | **Content Purpose** |
| Traceability Bases | load_basis | data/basis.json | basis.json | Loads the master list of source types (Basis) (e.g., GBIF, WoRMS). |
| Tags | populate_tags | N/A (Initialisation script) | | |
| Habitats | populate_habitats | N/A (Initialisation script) | | |

This is the bulk ingestion phase that operates in a loop (*for folder in data/groups/*/*), processing taxonomic data to ensure that all linked data are loaded together. In the case of the example provided, there is only one group (Amphibia), but more can be added as long as the same structure is maintained.

**Phase III: Loading of Taxonomic Data, Occurrences, and Genetic Information**

```
for folder in data/groups/*/
do
   # Taxonomy
   echo "$folder/taxonomy.csv"
   python manage.py load_taxonomy_new "$folder/taxonomy.csv"

   # Images
   echo "$folder/images.json"
   python manage.py load_images "$folder/images.json"

   # IUCN
   echo "$folder/iucn.json"
   python manage.py load_taxon_data "$folder/iucn.json"

   # Tags
   echo "$folder/tags.xlsx"
   python manage.py load_taxon_tags "$folder/tags.xlsx"
```

```
   # Occurrences
   for file in "$folder/occurrences/"*.json
   do
     echo "$file"
     python manage.py load_occurrences_new_synonyms "$file"
   done

    # Genetics
  for file in "$folder/genetics/"*.json
  do
    echo "$file"
    python manage.py load_occurrences_new_synonyms "$file"
  done
done

echo "Loading sources..."
python manage.py load_sources data/sources.csv
```

1. **Phase III-A (Taxonomy Load):**

   The *load_taxonomy_new* loads the complete taxonomy (*TaxonomicLevel*) and its correct execution is needed for the further steps. This script is executed first in each folder containing a CSV file (taxonomy.csv). Next, the *load_images*, *load_taxon_data*, and *load_taxon_tags* scripts are executed. These scripts read the scientific name from their input files and locate the internal ID of the reference taxon in the *TaxonomicLevel* table loaded previously. They then load the data associated with it, such as images, conservation status, or ecological data of interest (Tab. 14).

**Table 14.** Details of the commands, files, and paths where the files must be stored for uploading taxonomy and tags information.

| **III-A. Load by Taxonomic Group (Loop for *folder in data/groups/*/*)** | | | | |
|---|---|---|---|---|
| **Data loaded** | **Command (Django)** | **File path (from load_db.sh)** | **Sample File** | **Content Purpose** |
| Taxonomy | load_taxonomy_new | folder/taxonomy.csv | taxonomy.csv | Loads the complete taxonomic hierarchy (TaxonomicLevel and Authorship). |
| Images | load_images | folder/images.json | images.json | Loads image links linked to the taxonomy. |

| | | | | |
|---|---|---|---|---|
| IUCN / Taxon Data | load_taxon_data | folder/iucn.json | iucn.json | Loads IUCN conservation information (IUCNData). |
| Tags (Specific) | load_taxon_tags | folder/tags.xlsx | tags_amphibia.xlsx | Loads ecological attributes (e.g., System, Directive, HabitatTaxonomy). |

2. **Phase III-B (Occurrences and Genetics):**
   The *load_occurrences_new_synonyms* script is to process both JSON files from occurrences (e.g., *Alytes muletensis.json*) and genetic sequences (e.g., *Alytes muletensis_ncbi.json*). In both cases, the script looks up the scientific name to link the record to the correct taxonomy and, in the case of genetics, to link the *Sequence* to an existing or new *Occurrence* (Tab. 15).

**Table 15.** Details of the commands, files, and paths where the files must be stored for uploading occurrences and genetic data, using the *load_gadm* script.

| III-B. Occurrences and Genetic Information | | | | |
|---|---|---|---|---|
| **Data loaded** | **Command (Django)** | **File path (from load_db.sh)** | **Sample File** | **Content Purpose** |
| Occurrences | load_occurrences_new_synonyms | folder/occurrences/*.json | Alytes muletensis.json | Loads occurrences (location and date data) and genetic sequences associated with the previous taxonomy. |
| Genetics | load_occurrences_new_synonyms | folder/genetics/*.json | Alytes muletensis_ncbi.json | |

### S3.2. Foreign Key Resolution Logic

Once, the loading scripts are designed, they do not require the internal ID (TaxonomicLevel -> ID) in the input files (CSV/JSON), which simplifies data preparation (Tab. 16).

**Table 16.** Detail of the connection between tables using fields defined as Foreign Keys.

| **Table** | **FK Field** | **Requirement** | **Resolution Mechanism** |
|---|---|---|---|
| Occurrence | taxonomy | Scientific name (e.g., "Alytes muletensis") | Look up the scientific name in the TaxonomicLevel table (Tax.). If found, it associates the occurrence (Occ.) with its taxon. Occ. -> Tax. |

| | | | |
|---|---|---|---|
| Sequence | occurrence | Occurrence ID -> ID (Depends on the input JSON format) | Given that genetic metadata associates genetic information (Seq.) with a specific occurrence, they represent occurrence details according to the system architecture. The script must generate a temporary occurrence if it does not find one, and then link it to that occurrence and in turn to the target taxon. Seq. -> Occ. -> Tax. |
| TaxonTag | tag | Name of the tag (e.g., "established") | The script looks up the name field in the Tag table to establish the FK. |

#### S3.3. Automated Traceability Logic (versioning)

We integrate traceability management into the core of the framework to automate workflow and eliminate the need for users to manually define batches in the command line. Each time we run a load script (*load_*.py*), the system autonomously generates a new record in the *Batch* table with an initial status of 'PENDING'. As the script inserts new records into the target tables, whether taxonomic levels, occurrences, or authorships, they are automatically associated with the identifier of that newly created batch.

### S4. Technical API Documentation

We built the API system with the objective of centralising and standardising researchers' access to data and basic analysis services. We have versioned the main access point under the path */api/v1*, which guarantees stability and facilitates the future implementation of upcoming versions that can coexist. In addition, to make the API fully accessible, we provide comprehensive interactive documentation based on the OpenAPI (Swagger; drf-yasg, 2023) standard, available at */api/docs*.

In each of the following sections, we will define: the general endpoint structure (*/api/v1/[thematic_module]*), the table with the common query parameters (*name*, *taxonomy*, etc.) that we apply across all endpoints of that app, and finally, the table that lists the available endpoints along with the exclusive and necessary filtering parameters for each type of operation. Finally, we provided several examples of requests, and the expected outcomes. See Tables 17-28 for the detailed information.

### S4.1. General Endpoints and Documentation

Base Path:

*/api/*

**Table 17.** Description of the general endpoints into which the API is divided.

| Module | Endpoint (URL) | Description | Purpose |
|---|---|---|---|
| API | /status | Returns the current API status and version (e.g., 1.0.0). | Service monitoring and health check. |
| | /v1/[thematic_module] | Base endpoint to which queries are sent to obtain data from different types (/api/v1/taxonomy). | Returns data from the database. |
| Documentation | /docs/ | Interactive graphical interface for the OpenAPI documentation (Swagger UI). | Endpoint description and query testing. |

### S4.2. Taxonomy endpoints

Base Path:

*/api/v1/taxonomy*

**Table 18.** Common parameters for the Taxonomy thematic module endpoints.

| Parameter | Type | Description | Values |
|---|---|---|---|
| name | String | Taxon name to search for (supports partial matches if exact=false). | Free text |
| parent | Integer | ID of the parent taxon to filter the results. | TaxonomicLevel ID |
| rank | Integer | Taxonomic rank to filter by (e.g., 6 for SPECIES). | Values defined in TaxonomicLevel.RANK_CHOICES |
| accepted | Boolean | Filters whether the taxon is the accepted name or a synonym. | true or false |

**Table 19.** List of the different endpoints of the Taxonomy thematic module with their specific parameters and a description of the information they return.

| Endpoint (URL) | Parameters | Description |
|---|---|---|
| /search | name, rank, exact | Search by taxon name. |
| /list | name, parent, rank, accepted, page, page_size | Gets a paginated list of taxa with filters. |
| /list/count | Filters from /list | Returns the total count of taxa that meet the filters. |
| /list/csv | Filters from /list | Generates a streaming CSV file with the filtered list of taxa. |
| /taxon | id (required) | Detail of a taxon by ID (Accepts POST/PUT/DELETE if authenticated). |
| /taxon/parent | id (required) | Gets the immediate parent taxon. |
| /taxon/children | id (required), page, page_size | Gets the direct children of a taxon. |
| /taxon/children/count | id (required) | Counts the number of direct children of a taxon. |

| | | |
|---|---|---|
| /taxon/sisters | id (required), page, page_size | Gets the sister taxa of a given ID. |
| /taxon/descendants/count | id (required) | Counts all descendants of a taxon. |
| /taxon/composition | id (required) | Returns the complete lower taxonomic composition of a taxon (e.g., the number of species in a genus within a family). |
| /taxon/synonyms | id (required) | Returns the list of synonyms associated with an accepted taxon. |
| /taxon/source | id (required) | Lists the sources (OriginId) that contributed to the record of that taxon. |
| /taxon/checklist | id (required) | Generates a complete list of all descendant taxa in .CSV format. |
| /authorship | id (required) | Detail of the authorship record by ID. |

Examples:

1. Descendant count:

   Request:
   ```
   GET /api/v1/taxonomy/taxon/descendants/count?id=7
   ```

   Response:
   
   ```
   {
     "genus": 1,
     "species": 2
   }
   ```
   

2. Get lower taxonomy of the taxon:

   Request:
   ```
   GET /api/v1/taxonomy/taxon/composition?id=2
   ```

   Response:
   
   ```
   [
     {
       "id": 3,
       "name": "Chordata",
       "rank": 1,
       "totalSpecies": 5
     }
   ]
   ```
   

### S4.3. Occurrences endpoints

Base Path:

*/api/v1/occurrences*

**Table 20.** Common parameters for the Occurrences thematic module endpoints.

| Parameter | Type | Description | Values |
|---|---|---|---|
| taxonomy | Integer | Taxon ID. | TaxonomicLevel ID |
| voucher | String | ID of the physical sample voucher (e.g., herbarium number). | Free text |
| basisOfRecord | Integer | Record type (e.g., 4 for HUMAN_OBSERVATION). | Occurrence.BASIS_OF_RECORD values |
| year, month, day | Integer | Filter by collection date. | 1 to 12 (month), 1 to 31 (day), Year (>1900) |
| geographicalLocation | String | Name of a geographical unit to filter by spatial overlap. | Free text (e.g., "Mallorca") |

**Table 21.** List of the different endpoints of the Occurrences thematic module with their specific parameters for each request and a description of the information they return.

| Endpoint (URL) | Parameters | Description |
|---|---|---|
| / | id (required) | Detail of an occurrence. |
| /map | Filters from /list | Gets a list of occurrences with geospatial fields optimised for rendering. |
| /list | Filters from the table above, page, pageSize | Gets a paginated list of occurrences with complex filters. |
| /list/download | Filters from /list | Generates a streaming .CSV file with the filtered occurrences. |
| /list/count | Filters from /list | Counts the total number of occurrences that meet the filters. |
| /stats/month | taxonomy (required) | Returns the count of occurrences grouped by month for a taxon. |
| /stats/year | taxonomy (required) | Returns the count of occurrences grouped by year for a taxon. |
| /stats/source | taxonomy (optional) | Returns the count of occurrences grouped by their source (Basis). |

| | | |
|---|---|---|
| /stats/children | taxonomy (required) | Count of occurrences for a taxon and all its descendants. |

Examples:

1. Statistics by year:

   Request:

```
GET /api/v1/occurrences?id=4
```

   Response:



```
{
  "id": 4,
  "basisOfRecord": "human_observation",
  "coordinateUncertaintyInMeters": 28060,
  "decimalLatitude": 39.68952,
  "decimalLongitude": 2.88966,
  "day": 6,
  "depth": null,
  "elevation": null,
  "eventDate": "2024-05-06",
  "month": 5,
  "taxonomy": {
    "id": 14,
    "name": "Alytes muletensis",
    "taxonRank": "species",
    "scientificNameAuthorship": "(Sanchíz & Adrover, 1979)",
    "accepted": true,
    "acceptedModifier": "",
    "images": [
      {
        "source": {
          "id": 3,
          "basis": {
            "id": 5,
            "name": "iNaturalist",
            "acronym": ""
          },
          "dataType": "image",
          "extractionMethod": "api",
          "url":
"https://inaturalist-open-data.s3.amazonaws.com/photos/{id}"
        },
```

```
        "externalId": "61080851/medium.jpeg",
        "attribution": "(c) Gert Jan Verspui, some rights reserved
(CC BY-NC), uploaded by Gert Jan Verspui"
      }
    ],
    "parent": 10
  },
  "voucher": "215655910",
  "year": 2024,
  "sources": [
    {
      "source": {
        "id": 6,
        "basis": {
          "id": 1,
          "name": "Global Biodiversity Information Facility",
          "acronym": "GBIF"
        },
        "dataType": "occurrence",
        "extractionMethod": "api",
        "url": "https://www.gbif.org/occurrence/{id}"
      },
      "externalId": "4863852296",
      "attribution": null
    },
    {
      "source": {
        "id": 7,
        "basis": {
          "id": 1,
          "name": "Global Biodiversity Information Facility",
          "acronym": "GBIF"
        },
        "dataType": "dataset_key",
        "extractionMethod": "api",
        "url": "https://www.gbif.org/dataset/{id}"
      },
      "externalId": "50c9509d-22c7-4a22-a47d-8c48425ef4a7",
      "attribution": null
    }
  ],
  "location": {
    "id": 28,
    "parent": 4,
    "name": "Inca",
    "rank": "municipality"
```

```
  }
}
```

2. Count by source and taxon:

   Request:

```
GET /api/v1/occurrences/stats/source?taxonomy=14
```

   Response:

```
[
  {
    "source": "gbif",
    "count": 210
  }
]
```

### S4.4. Genetics endpoints

Base Path:

*/api/v1/genetics*

**Table 22.** Common parameters for the Genetics thematic module endpoints.

| Parameter | Type | Description | Values |
|---|---|---|---|
| taxonomy | Integer | Taxon ID to filter sequences or associated markers. | TaxonomicLevel ID |
| marker | Integer | Marker ID to filter sequences. | Marker ID |
| published_date_year | Integer | Filter by sequence publication year. | Year (>1990) |
| isolate | String | Filter by sequence isolation code. | Free text |

**Table 23.** List of the different endpoints of the Genetics thematic module with their specific parameters for each request and a description of the information they return.

| Endpoint (URL) | Parameters | Description |
|---|---|---|
| /marker | id (required) | Detail of a genetic marker. |

| | | |
|---|---|---|
| /marker/search | name (required), exact | Search by marker name. |
| /marker/list | name, product, is_relevant, page, pageSize | Gets a paginated list of markers. |
| /marker/list/count | Filters from /marker/list | Counts the total number of markers. |
| /sequence | id (required) | Detail of a DNA sequence. |
| /sequence/list | Filters from the table above, page, pageSize | Gets a paginated list of sequences. |
| /sequence/list/count | Filters from /sequence/list | Counts the total number of sequences. |
| /sequence/list/csv | Filters from /sequence/list | Generates a CSV with the filtered list of sequences. |
| /sequence/source/count | marker (optional), published_date_year | Count of sequences grouped by the source of origin (Basis). |
| /sequence/source/csv | Filters from /sequence/source/count | Downloads the result of the sequence count by source in CSV format. |

Examples:

1. Sequence count by source:

   Request:

   ```
   GET /api/v1/genetics/sequence/source/count?id=3&taxonomy=14
   ```

   Response:

   ```
   [
     {
       "source": "NCBI",
       "count": 1
     }
   ]
   ```

2. Sequence detail:

   Request:

   ```
   GET /api/v1/genetics/sequence?id=15
   ```

   Response:

```
{
  "id": 15,
  "sources": [
    {
      "source": {
        "id": 13,
        "basis": {
          "id": 3,
          "name": "National Center for Biotechnology Information",
          "acronym": "NCBI"
        },
        "dataType": "sequence",
        "extractionMethod": "api",
        "url": "https://www.ncbi.nlm.nih.gov/nuccore/{id}"
      },
      "externalId": "KJ858836.1",
      "attribution": null
    }
  ],
  "markers": [
    {
      "id": 9,
      "name": "ND2",
      "accepted": true
    }
  ],
  "occurrence": {
    "id": 824,
    "coordinateUncertaintyInMeters": null,
    "decimalLatitude": null,
    "decimalLongitude": null,
    "taxonomy": {
      "id": 14,
      "name": "Alytes muletensis",
      "taxonRank": "species"
    },
    "basisOfRecord": null,
    "depth": null,
    "elevation": null,
    "voucher": null
  },
  "isolate": "AM02",
  "definition": "Alytes muletensis isolate AM02 tRNA-Met gene, partial
sequence; and NADH dehydrogenase subunit 2 (ND2) gene, partial cds;
mitochondrial",
  "publishedDate": "2014-07-16"
}
```

### S4.5. Tags endpoints

Base Path:

*/api/v1/tags*

**Table 24.** Common parameters for the Tags thematic module endpoints.

| Parameter | Type | Description | Values |
|---|---|---|---|
| taxonomy | Integer | Taxon ID to get associated attributes. | TaxonomicLevel ID (required by all endpoints) |

**Table 28.** List of the different endpoints of the Tags thematic module with their specific parameters for each request and a description of the information they return.

| Endpoint (URL) | Parameters | Description |
|---|---|---|
| / | taxonomy (required) | List of generic tags (TaxonTag) associated with a taxon. |
| /directives | taxonomy (required) | Lists the status of a taxon under conservation directives (CEEA, LESPRE, Birds, Habitats, CITES, Berne). |
| /habitats | taxonomy (required) | Lists the habitat types (HabitatTaxonomy) associated with a taxon. |
| /iucn | taxonomy (required) | Lists the IUCN Red List status by taxon and region. |
| /system | taxonomy (required) | Lists the ecological system of the taxon (freshwater/marine/terrestrial). |

Examples:

1. Directives under which a taxon is registered:

   Request:

```
GET /api/v1/tags/directives?taxonomy=14
```

   Response:



```
{
  "sources": [
    {
      "source": {
        "id": 4,
        "basis": {
          "id": 9,
```

```
          "name": "Tommaso Cancellario",
          "acronym": ""
        },
        "dataType": "taxon_data",
        "extractionMethod": "expert",
        "url": ""
      },
      "externalId": null,
      "attribution": null
    }
  ],
  "cites": false,
  "ceea": true,
  "lespre": false,
  "directivaAves": false,
  "directivaHabitats": false
}
```


2. Associated habitats:

   Request:

```
GET /api/v1/tags/habitats?taxonomy=14
```

   Response:


```
[
  {
    "id": 5,
    "name": "wetlands (inland)",
    "sources": [
      {
        "source": {
          "id": 1,
          "basis": {
            "id": 4,
            "name": "International Union for Conservation of
Nature",
            "acronym": "IUCN"
          },
          "dataType": "taxon_data",
          "extractionMethod": "api",
          "url": ""
        },
        "externalId": "5",
        "attribution": null
      }
```

```
    ]
  },
  {
    "id": 15,
    "name": "artificial/aquatic",
    "sources": [
      {
        "source": {
          "id": 1,
          "basis": {
            "id": 4,
            "name": "International Union for Conservation of
Nature",
            "acronym": "IUCN"
          },
          "dataType": "taxon_data",
          "extractionMethod": "api",
          "url": ""
        },
        "externalId": "15",
        "attribution": null
      }
    ]
  },
  {
    "id": 14,
    "name": "artificial/terrestrial",
    "sources": [
      {
        "source": {
          "id": 1,
          "basis": {
            "id": 4,
            "name": "International Union for Conservation of
Nature",
            "acronym": "IUCN"
          },
          "dataType": "taxon_data",
          "extractionMethod": "api",
          "url": ""
        },
        "externalId": "14",
        "attribution": null
      }
    ]
  }
```

```
]
```

### S4.6. Geography endpoints

Base Path:

*/api/v1/geography*

**Table 25.** Common parameters for the Geography thematic module endpoints.

| Parameter | Type | Description | Values |
|---|---|---|---|
| id | Integer | Geographical level ID. | GeographicLevel ID |
| name | String | Name of the geographical level (e.g., "Mallorca"). | Free text |
| rank | Integer | Geographical rank (e.g., 4 for MUNICIPALITY). | GeographicLevel.RANK_CHOICES values |

**Table 26.** List of the different endpoints of the Geography thematic module with their specific parameters for each request and a description of the information they return.

| Endpoint (URL) | Parameters | Description |
|---|---|---|
| /level | id (required) | Detail of a geographical level. |
| /level/parent | id (required) | Gets the ancestral geographical levels of an ID. |
| /level/children | id (required), page, pageSize | Gets the direct children of a geographical level. |
| /search | name (required), exact | Search for geographical levels by name. |

Examples:

1. Get geographical children:

   Request:

   ```
   GET /api/v1/geography/level/children?id=1
   ```

   Response:

   ```
   [
     {
   ```

```
    "id": 2,
    "parent": 1,
    "name": "Eivissa",
    "rank": "island",
    "decimalLatitude": 38.98372,
    "decimalLongitude": 1.40618,
    "coordinateUncertaintyInMeters": 24366,
    "area": “{...}”
  }
]
```

2. Search for geography by name:

   Request:

```
GET /api/v1/geography/search?name=menor&exact=false
```

   Response:

```
[
  {
    "id": 5,
    "parent": 1,
    "name": "Menorca",
    "rank": "island"
  },
  {
    "id": 45,
    "parent": 5,
    "name": "Ciutadella de Menorca",
    "rank": "municipality"
  }
]
```

### S4.7. Versioning endpoints

Base Path:

*/api/v1/versioning*

**Table 27.** List of the different endpoints of the Versioning thematic module with their specific parameters for each request and a description of the information they return.

| **Endpoint (URL)** | **Parameters** | **Description** |
|---|---|---|
| /basis | id (required) | Detail of a source type (Basis). |

| | | |
|---|---|---|
| /basis/list | page, pageSize | Lists the available source types (Basis). |
| /basis/statistics | basis (optional) | Returns the contribution statistics for each Source grouped by its Basis. |
| /basis/list/count | name, exact | Counts the total number of source types. |
| /basis/search | name (required), exact | Search by source type name. |
| /source | id (required) | Detail of a specific source. |
| /source/list | basis, data_type, page, pageSize | Lists the specific sources (Source). |
| /source/list/count | Filters from /source/list | Counts the total number of specific sources. |
| /origin | id (required) | Detail of an origin identifier (OriginId). |

Examples:

1. Contribution statistics by source:

   Request:

   ```
   GET /api/v1/versioning/basis/statistics?id=1
   ```

   Response:

   ```
   [
     {
       "id": 6,
       "dataType": "occurrence",
       "extractionMethod": "api",
       "count": 809
     },
     {
       "id": 7,
       "dataType": "dataset_key",
       "extractionMethod": "api",
       "count": 16
     },
     {
       "id": 5,
       "dataType": "taxon",
       "extractionMethod": "api",
       "count": 0
     }
   ]
   ```

2. Detail of a data source:

Request:

```
GET /api/v1/versioning/basis?id=1
```

Response (simplified JSON):


```
{
  "id": 1,
  "name": "Global Biodiversity Information Facility",
  "acronym": "GBIF",
  "type": "database",
  "url": "https://www.gbif.org/",
  "description": "GBIF—the Global Biodiversity Information
Facility—is an international network and data infrastructure
funded by the world's governments and aimed at providing anyone,
anywhere, open access to data about all types of life on Earth.",
  "authors": "['Gray', 'Wagler']",
  "citation": "GBIF.org (2024), GBIF Home Page. Available from:
https://www.gbif.org",
  "contact": "",
  "image": null
}
```


### S5. Front-end deployment

In this section, we detail the architecture of the web interface (front-end). We have built the system on Next.js 15 (Vercel Inc., 2025; React 19 - React Team, 2024), adopting the App Router architecture to maximise performance through hybrid rendering. This decision allows us to decouple the presentation logic from the data logic, ensuring the long-term scalability and maintainability of the project. The front-end is connected to the back-end *via* the RESTful API system.

The first step to build up the front-end is to clone the repository from GitHub:

```
git clone
https://github.com/centrebalearbiodiversitat/BiOS_frontend.git
```

Access the project directory:

```
cd BiOS_next
```

Installing dependencies: Install the necessary libraries defined in *package.json*.

```
npm install
```

To configure the API connection, create a *.env* file by duplicating the *.env.template,* and renaming it to *.env,* in the front-end root to indicate where the back-end is listening (by default on port 8000 locally). Setting DEBUG will run the Next.js server in debug mode. The MAINTENANCE flag disables the entire user interface by informing with a message that maintenance is in progress.

```
# .env
DEBUG=true
MAINTENANCE=false
API_BASE_URL="http://localhost:8000"
API_PATH="/api/v1"
```

**Running in development server mode**

To start the development server mode with hot reload:

```
npm run dev
```

**Deploy in production mode**

To deploy to a production server, generate the optimised version and run the server.

**NOTE:** Make sure to configure the *.env* file properly before deploying to production.

```
npm run build
npm start
```

The web application will be available at *http://localhost:3000*.

To confirm that both systems are connected correctly, access *http://localhost:3000/en/map* (or the configured port).
If the MapLibre base map loads correctly and the side filters display taxonomic options, the connection to the API is correct.
